# Low dimensional dynamics for working memory and time encoding

**DOI:** 10.1101/504936

**Authors:** Christopher J. Cueva, Alex Saez, Encarni Marcos, Aldo Genovesio, Mehrdad Jazayeri, Ranulfo Romo, C. Daniel Salzman, Michael N. Shadlen, Stefano Fusi

## Abstract

Our decisions often depend on multiple sensory experiences separated by time delays. The brain can remember these experiences and, simultaneously, estimate the timing between events. To understand the mechanisms underlying working memory and time encoding we analyze neural activity recorded during delays in four experiments on non-human primates. To disambiguate potential mechanisms, we propose two analyses, namely, decoding the passage of time from neural data, and computing the cumulative dimensionality of the neural trajectory over time. Time can be decoded with high precision in tasks where timing information is relevant and with lower precision when irrelevant for performing the task. Neural trajectories are always observed to be low dimensional. These constraints rule out working memory models that rely on constant, sustained activity, and neural networks with high dimensional trajectories, like reservoir networks. Instead, recurrent networks trained with backpropagation capture the time encoding properties and the dimensionality observed in the data.

## 1 Introduction

Humans and other animals are free from the immediacy of reflex actions thanks to their ability to preserve information about their sensory experiences. When events like sensory inputs, decisions or motor responses are separated by time delays, the subject has to be able to propagate information across these delays (e.g. the identity of a visual stimulus). Moreover, it is often the case that the duration of the delay intervals can be fundamental for interpreting incoming streams of sensory stimuli, requiring the subject to measure the time that passes between one relevant event and the next. The ability to propagate in time the information about the event preceding the delay relies on what is often defined as working memory, and it has been extensively studied^1, 2^. Analogously, there are several studies on the capacity of humans and animals to measure the time that has elapsed since the event^3, 4^. Here we analyzed the delay activity recorded in monkeys during four different experiments to understand the dynamics of the neural mechanisms that enable monkeys to preserve over time the information about a particular event and, at the same time, to measure the interval that passed since that event. We considered three classes of mechanisms that have been suggested by previous theoretical work (Figure 1).

**Figure 1:**
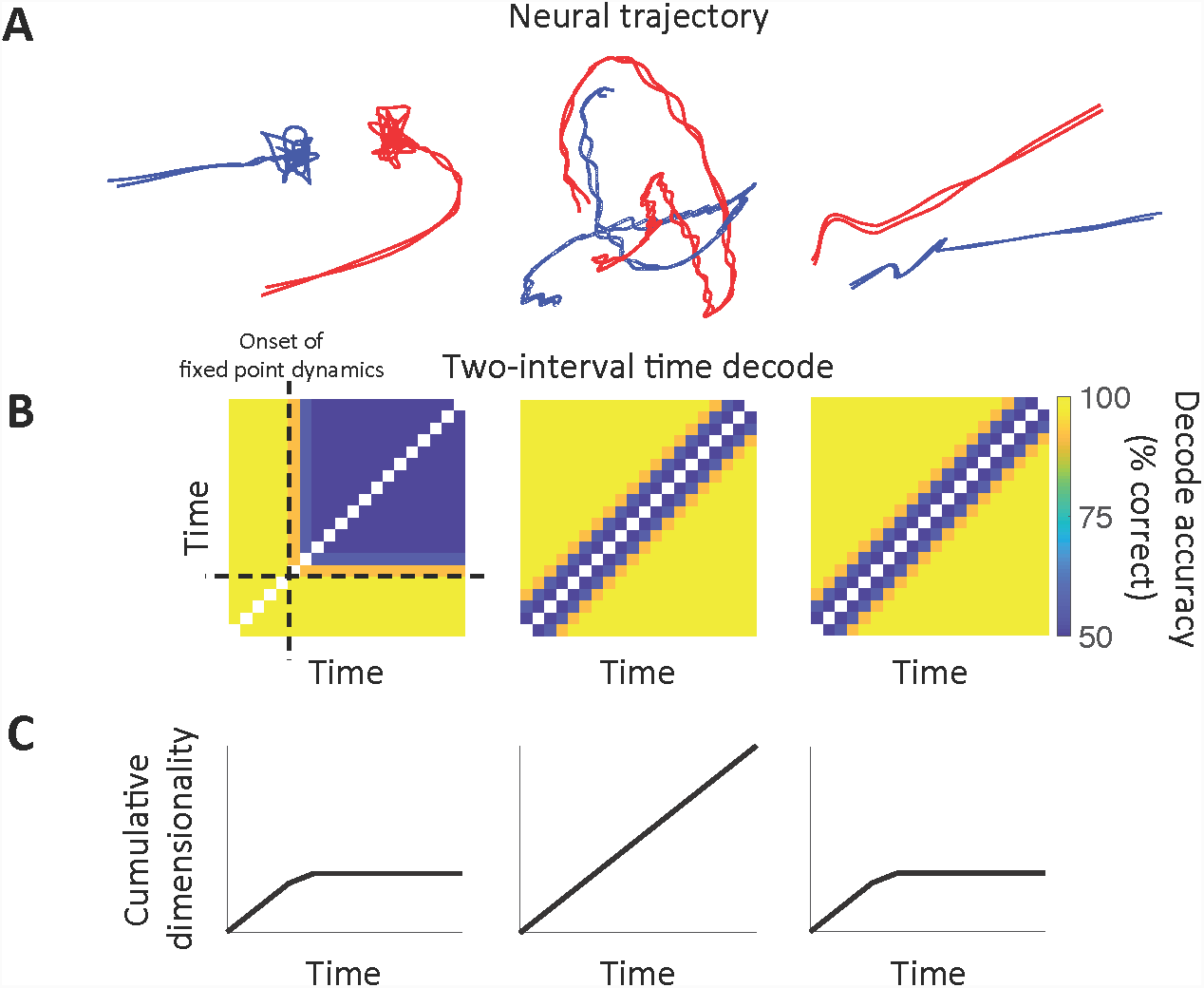
Three different types of neural dynamics, which can be identified by decoding time and dimensionality. **(A)** Trajectories in the firing rate space: the firing rates of a population of simulated neurons are shown after they have been projected onto a two dimensional space capturing the largest variance in their trajectories over time. On the left, a transient response is followed by attractor dynamics. Information about two behavioral states is stored in separate fixed points colored in red and blue. The two lines for each behavioral state correspond to two different trials. These fixed points are attractors of the dynamics and the fluctuations around them are due to noise. In the center, a randomly connected “reservoir” of neurons generates chaotic trajectories. The trajectories have been stabilized as in Laje and Buonomano^10^. The neural activity at each timepoint is unique and these changing firing rates can be used as a clock to perform different computations at different times. Importantly, the red and blue trajectories are distinct and linearly separable for all times, so also the behavioral state is encoded throughout the interval, as in the attractor dynamics. On the right: low dimensional trajectories a transient is followed by linearly ramping neural responses. **(B)** Decoding time from the data shown in (A). Neural firing rates are constant at a fixed point, and a classifier cannot discriminate different timepoints. In contrast, in the reservoir computing framework the neural activity at each timepoint is unique and it is possible to decode the ‘passage of time’ from the neural population. Decoding time from neural activity (*Methods* Figure M1) helps identify the contrasting neural regimes. Pixel(i,j) is the decode accuracy of a binary classifier trained to discriminate timepoints i and j. On the left, the block of time where the decode is near chance level (50%) is a signature of fixed point dynamics. On the right and center, it is possible to decode time (down to some limiting precision due to noise in the firing rates) but these two dynamical regimes, while different than the fixed point dynamics, yield the same two-interval time decode. However, they are disambiguated by computing the cumulative dimensionality over time. **(C)** The cumulative dimensionality of the neural activity over time increases linearly in the standard stabilized reservoir network (center). This is in contrast to fixed point and ramping dynamics where the cumulative dimensionality increases during an initial transient and then plateaus during the fixed point and ramping intervals.

The first mechanism is often used to model working memory (see e.g. 5, 6) and it is based on the hypothesis that there are neural circuits that behave like an attractor neural network^5, 7, 8^ (Figure 1, left column), in which different events (e.g. different sensory stimuli) lead to different stable fixed points of the neural dynamics. Persistent activity, widely observed in many cortical areas, has been interpreted as an expression of this attractor dynamics (see e.g. 9). For these dynamical systems, the information about the event preceding the delay is preserved as long as the neural activity remains in the vicinity of the fixed point representing the event. However, once the fixed point is reached, the variations of the neural activity are only due to noise; all timing information is lost and time is not encoded.

Time and memory encoding can be obtained simultaneously in a category of models known as reservoir networks, liquid state machines or echo state networks^11–13^. These are recurrent neural networks (RNNs) with random connectivity that can generate high dimensional chaotic trajectories (Figure 1A, center). If these trajectories are reproducible, then they can be used as clocks as the network state will always be at the same location in the firing rate space after a certain time interval. Thanks to the high dimensionality, one can implement the clock using a simple linear readout. Moreover, a linear readout is also sufficient to decode any other variable that is encoded in the initial state. To identify this computational regime we note that a prediction of the reservoir computing framework is that neural activity at each timepoint is unique. If this is true, then it will be possible to decode the ‘passage of time’ from the neural population (Figure 1B center and *Methods* Figure M1), regardless of whether timing information is relevant for the task or not. However, there are a few problems with these models. In principle they are very powerful, as they can generate any input-output function (output function of spatiotemporal inputs). However, this would require an exponential number of neurons, or equivalently, the memory span would grow only logarithmically with the number of neurons. Moreover, the trajectories are chaotic, and so inherently unstable and not robust to noise. Although, recent theoretical work^10, 14^ has demonstrated ways of making them robust.

The third category of models is one in which the activity varies in time but across trajectories that are low dimensional. For these models, it is still possible to encode time and also to encode different values of other variables along separate trajectories (Figure 1, right column). Working memory, in these models, does not rely on constant rates around a fixed activity pattern as in standard attractor models, however, the low-dimensional evolving trajectories can still provide a substrate for stable memories, similar to attractor models of memory storage via constant activity. One advantage of a low-dimensional evolving trajectory is that time can also be encoded, something not possible with constant neural activity.

Here we show that the last scenario is compatible with four datasets from monkeys performing a diverse set of working memory tasks. Time can be decoded with low precision in tasks where timing information is irrelevant, if one excludes from the analysis the short time interval immediately following task relevant events (e.g. the offset of the visual stimulus) which, most likely, provides an external clock. Time can be decoded with higher precision in tasks where it is relevant, consistent with the idea that stable neural trajectories act as a clock to perform the task. However, neural trajectories for all tasks are low dimensional; they evolve on slow timescales such that the cumulative dimensionality of the neural activity over time is low.

Interestingly, many of the observed features of the data can be reproduced using recurrent neural network models trained to perform the experimental tasks using backpropagation through time.

## 2 Decoding time from neural data

Decoding time from the recorded patterns of activity is a powerful way of gaining insight into the dynamics of the neural circuits. Indeed, temporal information can be extracted only if some of the components of the neural dynamics are reproducible. In other words, the two-interval time decode analysis can help us to identify the time variations that are consistent across trials and to focus on the actual dynamics of the neural circuits, ignoring the often large dynamical components that are just noise.

To assess whether there is any information about the time that passed since the last sensory event, we first train a decoder to discriminate between two different time intervals (Figure 1. This type of discriminability is a necessary condition for time to be encoded in the neural activity (if all time intervals are indistinguishable, then of course time is not encoded). This preliminary analysis also reveals that time is encoded in different ways in the different time intervals.

We start with the trace-conditioning experiment conducted by Saez et al.^15^ but we will later report the results of the same analysis for all datasets. In Saez et al.^15^ monkeys were presented with one of two visual stimuli, A or B. After a 1.5 second delay period the monkey was either rewarded or not. This is a context dependent task: in context 1, stimulus A is rewarded and stimulus B is not, whereas, in context 2, the associations are reversed (stimulus A is not rewarded and stimulus B is rewarded). The trials are presented in contextual blocks; all trials within a block have the same context. The monkey displays anticipatory behavior and in context 1 starts licking the water spout after stimulus A and not after stimulus B. In context 2 the monkey also performs as expected, licking after stimulus B and not after stimulus A. In Saez et al.^15^ it was shown that the monkey is not just relearning the changing associations between stimuli and reward but has actually created an abstract representation of context (see also Bernardi et al.^16^).

The two-interval time decode analysis is shown in Figure 2 for three brain areas: the orbitofrontal cortex (OFC), the anterior cingulate cortex (ACC) and the amygdala. We discretized time (100 ms bins) and trained a binary classifier to discriminate between the patterns of activity recorded at two different discrete times. We then plotted the accuracy of the classifier for every pair of time intervals, constructing the matrices of Figures 2A and 2C (for more details see *Methods* Figure M1). In Figure 2A we considered the delay interval between the offset of the visual stimulus and before the reward, and in Figure 2C the interval preceding the presentation of the visual stimulus. The decode accuracy is near 100% during the initial *∼*500 ms after the offset of the visual stimulus (Figure 2A) and during the presentation of the visual stimulus (Figure 2C), but it decreases near chance level for the remainder of the delay period and in the interval preceding the visual stimulus. This is observed in all three brain areas. The blocks of time where the two-interval time decode is near chance level (50%) are consistent with constant firing patterns fixed points of the neural dynamics; if the neural activity at all timepoints is similar it will not be possible to discriminate between different time intervals, and hence decode the passage of time. This is consistent with previous findings during working memory tasks, where constant firing patterns are displayed by the self-sustaining reverberations of activity hypothesized to support time invariant storage ^17–20^. Importantly, the inability to decode time from the neural data is not simply due to excessive noise as other task relevant variables, like context and whether the monkey receives a water reward or not, can be decoded as shown in Figure 2B and 2D. All these quantities could be decoded throughout the delay in all three brain areas, as already reported in Saez et al.^15^

**Figure 2:**
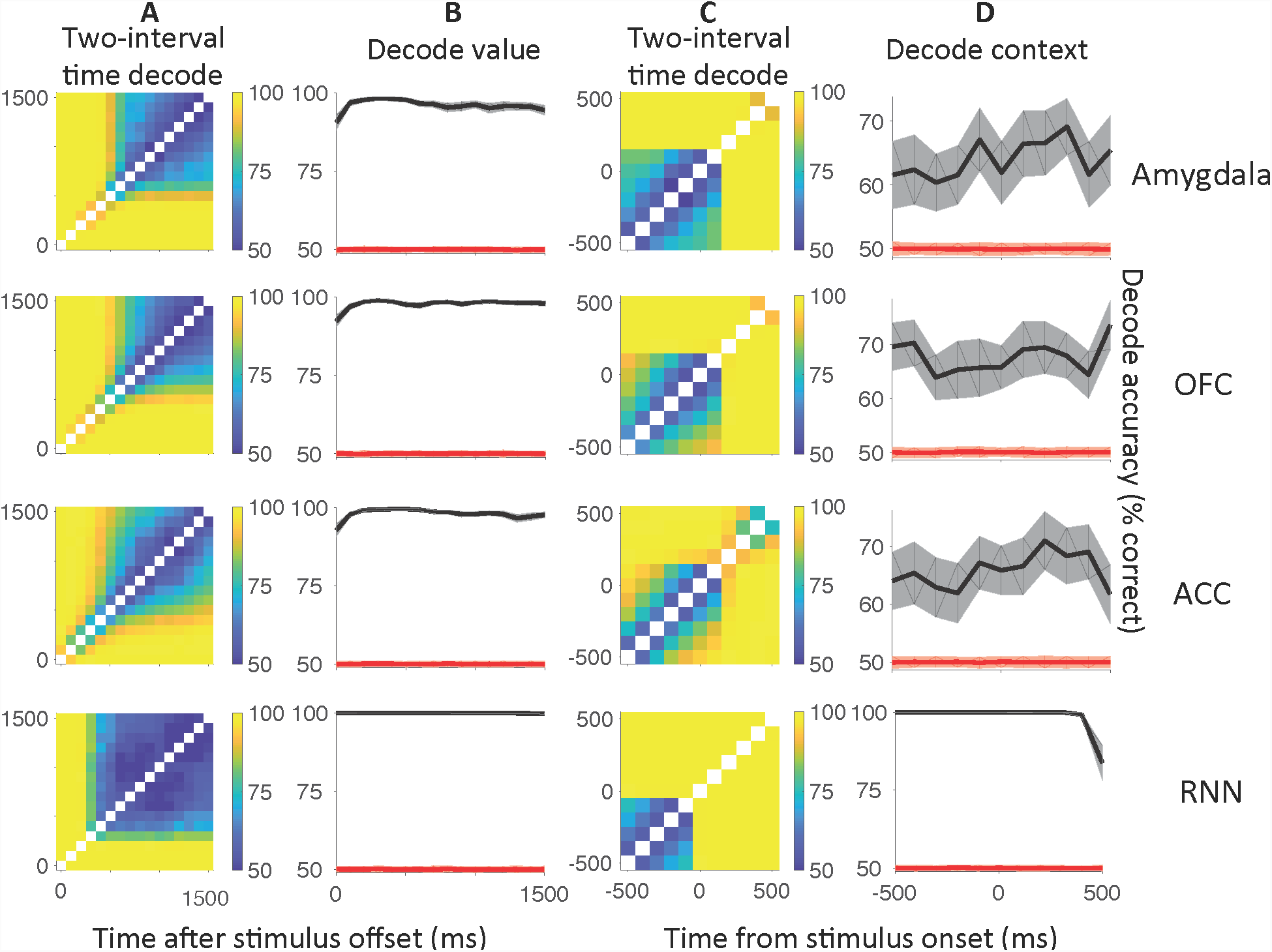
Decoding time from neural activity reveals signatures of fixed point dynamics. **(A, C)** Pixel(i,j) is the decode accuracy of a binary classifier trained to discriminate timepoints i and j using 100 ms bins of neural activity. The blocks of time where the decode is near chance level (50%) are signatures of fixed point dynamics. The pattern of fixed points seen in the data agree with the RNN model. In the model, the fixed points before stimulus onset store contextual information (column C) and the fixed points after stimulus offset encode the expected reward and the stimulus (column A). **(B, D)** Importantly, a linear classifier can easily discriminate other task relevant quantities during these time intervals so the poor time-decode is not simply due to noisy neural responses. The black curve shows the decode accuracy of a binary classifier trained to discriminate reward and no-reward trials (B) and trials from context 1 and context 2 (D) using 100 ms bins of neural activity. Error bars show two standard deviations.

### Reproducing the data with a recurrent neural network model

An artificial recurrent neural network (RNN) trained to reproduce monkey behavior on this task shows the same pattern of fixed point dynamics (Figure 2A and 2C, bottom row) and the two-interval time decode analysis produces results that are similar to those observed in the experiment. This is significant because neural connectivity in the artificial RNN was initialized randomly before training and unit activity was not constrained to replicate neural data during training; the artificial RNN was only told ‘what’ to do but not ‘how’ it should be done. The inputs to the RNN are time-varying signals representing experimental stimuli and we ‘train’ the RNN so its outputs are time-varying signals representing behavioral responses (anticipatory licking behavior). Importantly, during the delay there is no input and the dynamics are entirely driven by the recurrent dynamics. The training procedure adjusts the connection weights between units using backpropagation through time, as in pioneering work by Zipser et al.^8^, in which they also constructed a neural network that could reproduce delay activity data. The weights are tuned so that every input pattern produces the desired output pattern. In our case, as the tasks are significantly more complex than those modeled in Zipser et al.^8^ weights were optimized with truncated Newton methods using backpropagation through time^21^. After training is complete the weights are fixed and the model produces the appropriate context dependent responses for any sequence of stimuli and changing contexts. The RNN is not explicitly given contextual information and must infer it from the pairing of stimulus and reward. In Figure 2A the fixed point dynamics appearing after stimulus offset in the RNN model correspond to the network entering a state of either reward expectation or no-reward expectation and may correspond to the monkey’s state of either licking in anticipation of reward or not-licking. The putative fixed point dynamics seen in the electrode data before stimulus onset (Figure 2C) are also present in the RNN model. The RNN model transitions to one of two fixed points during the intertrial interval to store contextual information between trials. This is surprising because we started with a randomly connected network that knew nothing about context or anything else; context was not present at the beginning of training and this information is never explicitly given to the network. The RNN formed an abstract understanding of the environment just by learning to generate the right behavior. This contextually dependent behavior is enabled by our use of a recurrent neural network (see also Figure M7 for an extended explanation).

## 3 Fixed points or slowly moving points?

The analysis of the activity recorded during the trace conditioning task appears to be compatible with fixed point dynamics. However, the delay between external events is only 1.5 seconds. So there is the possibility that the dynamics have variations on a longer time-scale that we cannot observe with this dataset. To explore this possibility, we next looked at a dataset with a longer interval between external events, namely, recordings from PFC in a vibrotactile discrimination task^22^ which has been extensively analyzed (see e.g. 23–27) and modeled (see e.g. 26–28). In this task a mechanical probe vibrates the monkey’s finger at one of 7 frequencies. Then there is either a 3 or 6 second delay interval before the monkey’s finger is vibrated again at a different frequency. The monkey’s task is to report whether the frequency of the second stimulus is higher or lower than that of the first. This dataset has already been analyzed in multiple ways and it is known that several neurons exhibit a time dependent ramping average activity^23, 24, 27^. However, time has never been explicitly decoded.

The longer delay period intervals used in this task reveal that it is possible to discriminate time intervals, but only if they are sufficiently separated, which means that time is encoded with a limited precision. Figure 3A shows the two-interval time decode analysis from PFC for delay intervals of 3 and 6 seconds. After an initial visual transient, whose duration is similar to the one observed in the dataset of Saez et al.^15^, the two-interval time decode accuracy decreases in a band around the diagonal. However, time intervals that are separated by more than 1 second can still be distinguished, indicating that time can actually be decoded but with a low precision. The precision is similar in the 3 and 6 seconds cases. Moreover, the two-interval time decode plots are similar to those of Saez et al.^15^ when one focuses on the initial part of the interval (supplementary Figure S1).

**Figure 3:**
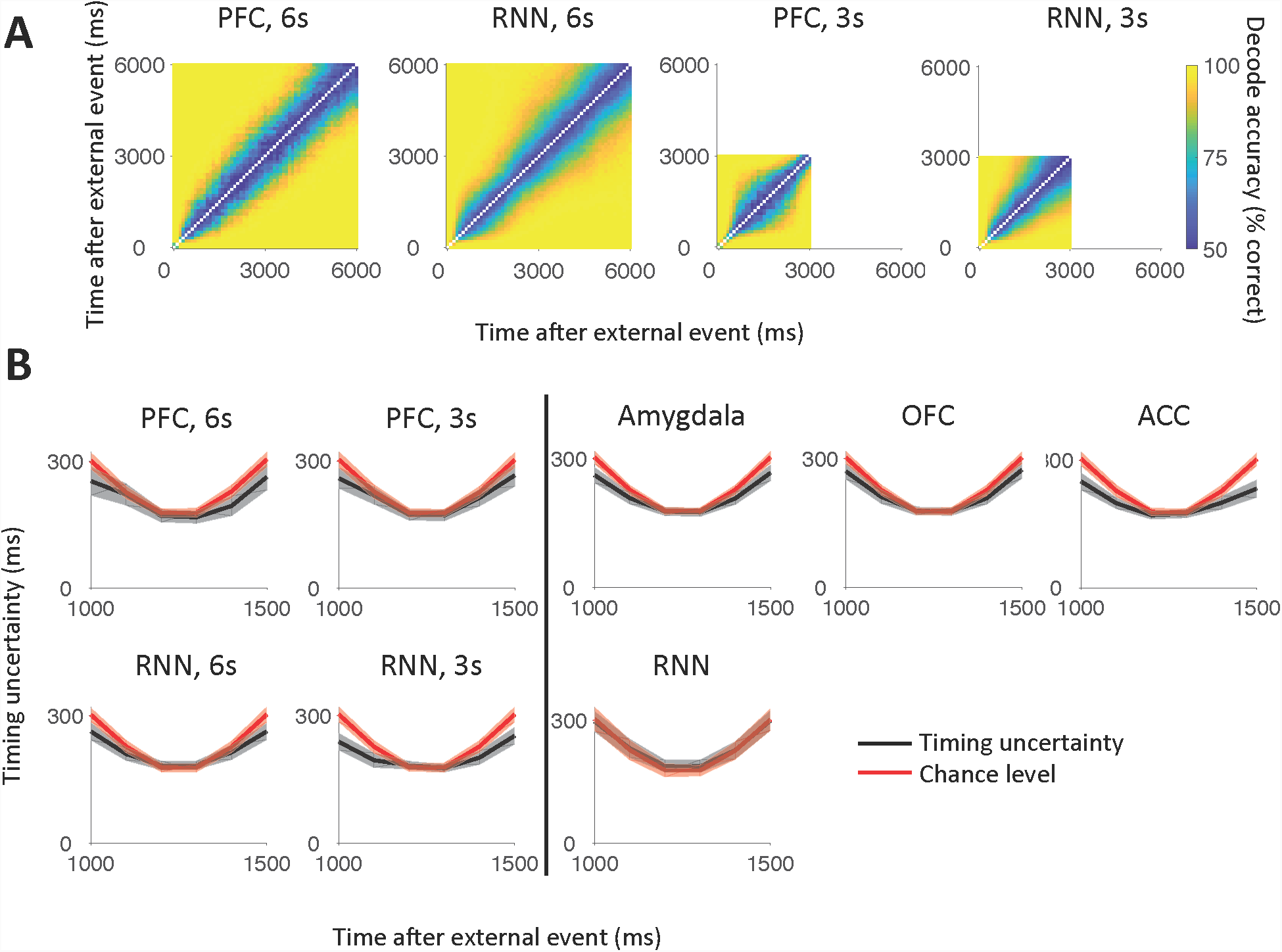
Network dynamics appear to evolve on a timescale of approximately half a second. **(A)** The two-interval time decode analysis from PFC for delay intervals of 3 and 6 seconds is shown on the right and left for both the data and RNN model. In each half second interval the decode accuracy is near chance level, however, the longer delay period intervals used in this task reveal neural dynamics that evolve over longer time scales. **(B)** The slowly varying dynamics observed in PFC are consistent with the putative fixed point dynamics observed in the Amygdala, OFC, and ACC. To quantify the temporal uncertainty at each point in time we train a classifier to estimate the timepoint a neural recording was made and then compare this prediction to the actual time; we repeat this classification for many trials obtaining a distribution about the true time. The ‘timing uncertainty’ is the standard deviation of this distribution and is shown in black. The chance level is shown in red. The timing uncertainty is shown during the 500 ms interval with putative fixed point dynamics in Amygdala, OFC, and ACC, and for an arbitrary 500 ms interval after the transient at the beginning of the trial in PFC. In all brain regions the timing uncertainty is near chance level. Longer time-scale fluctuations may not be as apparent in the Amygdala, OFC, and ACC because a shorter delay period was used in the experiment, preventing neural dynamics from evolving sufficiently in the absence of external stimuli. Error bars show two standard deviations.

The dynamics observed in PFC slowly evolve on a timescale of approximately half a second and are consistent with the putative fixed point dynamics observed in the Amygdala, OFC, and ACC (Figure 3B). In Figure 3B we see that, after the transient following stimulus offset, the timing uncertainty of the neural data is at chance level over an interval of half a second for both the PFC data of Romo et al.^22^ and the data of Saez et al.^15^

To quantify the timing uncertainty of the neural data at a given point in time we train a classifier to predict the time this recording was made, and then use the spread of predictions when classifying firing rates from different trials as our measure of timing uncertainty. This classifier takes the firing rates from all neurons at a given timepoint (time is discretized in 100 ms bins) and predicts the time this recording was made. This is in contrast to the two-interval time decode analysis where the binary classifier only discriminates between two timepoints; the classifier attempts to predict the actual timepoint within the trial, e.g. 1000 ms after stimulus offset, or, in other words, it decides what is the most likely class among all the classes that correspond to different time bins. For this reason it is a multi-class classifier.

The prediction of the multi-class classifier is compared to the actual time to obtain the timing uncertainty. After performing this classification on many trials we obtain a distribution of predictions around the true value (*Methods* Figure M2). The timing uncertainty shown in Figure 3B (black curves) is the standard deviation of this distribution of predicted values minus the true value. The chance level for the timing uncertainty (red curves in Figure 3B) is computed by training and testing the classifier on neural data with random time labels. The chance level is U-shaped as a classifier with uniform, random predictions can make larger errors when the true value is at the edge of the interval. In Figure 3B, the timing uncertainty is shown during the 500 ms interval with putative fixed point dynamics in Amygdala, OFC, and ACC, and for an arbitrary 500 ms interval after the transient at the beginning of the trial in PFC. In all brain regions the timing uncertainty is near chance level when this short interval is considered and plots are similar, even if the tasks and the brain areas are different.

Similar long timescale dynamics are generated by a neural network model trained to reproduce the experimentally observed behavior of discriminating frequency pairs, plus an extra anticipatory output that predicts the time of the next event after the delay period, namely, the delivery of the second vibrotactile frequency. The anticipatory output is essential; without it the RNN model uses only fixed point dynamics to store the frequency of the first stimulus and does not generate evolving dynamics. The network reproduces both the two-interval time decode plots of Figure 3A and the more quantitative analysis of Figure 3B.

## 4 Encoding time in tasks where timing is important

In both tasks we have considered the monkey was not explicitly required to keep track of timing information. In the randomly connected recurrent networks, with stabilized trajectories, proposed in Laje and Buonomano^10^ it should be possible to decode time whether the timing information is relevant for the task or not. However, it is also possible that the task actually shapes the necessary neural dynamics and time is encoded only when necessary. We analyzed two datasets in which timing information was necessary in order to solve the task. The ready-set-go interval reproduction task from Jazayeri and Shadlen^29^ required the monkey to keep track of the interval duration between the ready and set cues (demarcated by two peripheral flashes) in order to reproduce the same interval with a self initiated saccadic eye movement at the appropriate time after the set cue. The duration-discrimination task from Genovesio et al.^30^ required the monkey to compare the duration of two visual stimuli (S1 and S2) sequentially presented and then report which stimuli lasted longer on that trial. Each of the two stimuli could be either a red square or a blue circle.

We decoded the passage of time in these datasets during intervals in which the monkey had to keep track of timing information, i.e. the interval between ready and set cues for Jazayeri and Shadlen’s LIP data, and the S1 interval for Genovesio et al.’s PFC data. To see the neural dynamics evolve in the absence of external events we only included trials with over 1000 ms between external events. We found that we could decode time with higher precision than in the datasets where timing information was not explicitly required (Figure 4). These results provide support for the scenario in which the task can shape the neural dynamics depending on whether timing information is important or not in the task. In the case of the vibrotactile task analyzed in the previous section, time could be decoded with lower precision. One could argue that in that case the timing information is not strictly necessary to perform the task, but it could help to prepare the monkey for the arrival of the stimulus. So the difference between the three tasks in which we could decode time is in the relative importance of the timing information, which also seems to shape the neural dynamics.

**Figure 4:**
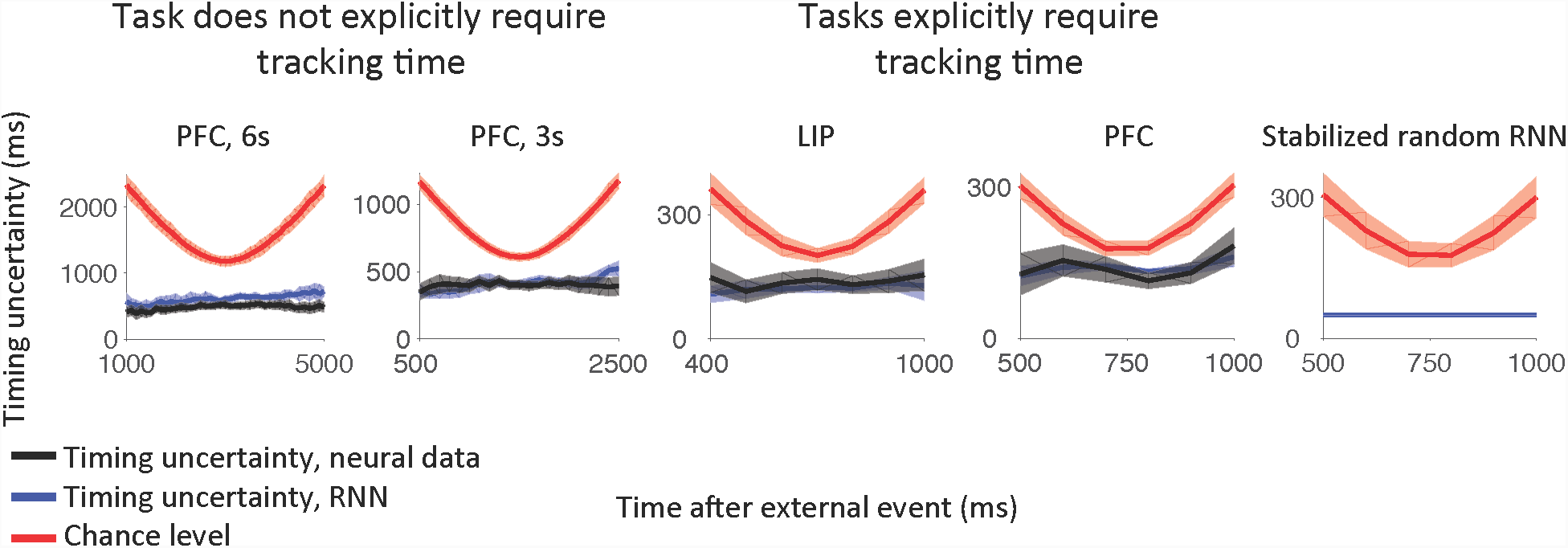
The timing uncertainty for the neural data (black curves) and RNN models (blue curves) is less than chance level (red curves) and has better resolution for tasks in which timing information is explicitly required, as in the duration-discrimination task of Genovesio et al.^30^ (PFC data) and the ready-set-go interval reproduction task of Jazayeri and Shadlen^29^ (LIP data). This is consistent with the idea that stable neural trajectories act as a clock to perform the task. Error bars show two standard deviations.

### The importance of ramping activity

Next we tried to identify the component of the dynamics that is most important for encoding time in all the cases in which we could decode it. The neural dynamics appear to be driven by the linear ‘ramping’ component of each neuron’s firing rate. Figure 5 shows the timing uncertainty in our ability to classify trials after the linear component is removed for all the experiments in which time could be decoded (compare to Figure 4). For each neuron, we calculated the linear fit to the average firing rate across trials, during the intervals shown in Figure 5. We then subtracted this linear fit from the neuron’s firing rate on every single trial. The mean firing rate across trials from two example neurons is shown in Figure 5A, before and after the linear fit is subtracted. After the linear ramping component is removed we compute the timing uncertainty, by classifying single trials as we do to compute the timing uncertainty in all figures, and see that the timing uncertainty is near chance level for both tasks in which timing information is, and is not, explicitly relevant (Figure 5B, black curves). This is also observed in the RNN models we trained to solve the experimental tasks (Figure 5B, blue curves). In contrast, for the stabilized random RNN^10^ it is still possible to decode time with high accuracy even after the linear component has been removed (Figure 5B, rightmost panel).

**Figure 5:**
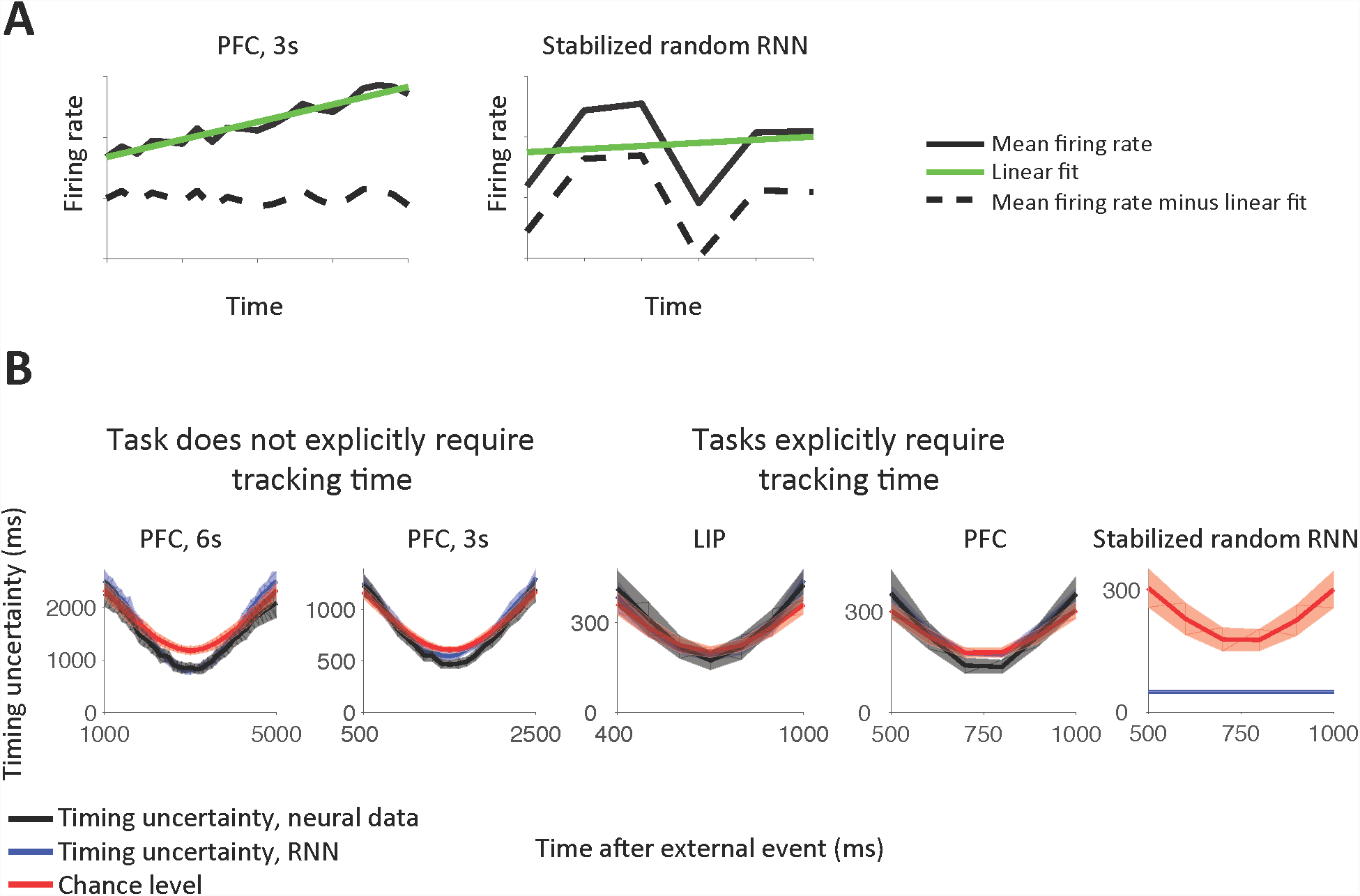
The linear “ramping” component of each neuron’s firing rate drives the decoder’s ability to estimate the passage of time (compare with Figure 4). **(A)** For each neuron, we calculate the linear fit to the average firing rate across trials during the same time interval as in Figure 4. We then subtract this linear fit from the firing rate on every single trial and calculate the timing uncertainty. **(B)** After the linear ramping component is removed the timing uncertainty in both the neural data (black curves) and trained RNN models (blue curves) is near chance level (red curves). In contrast, for the untrained RNN with stabilized chaotic dynamics^10^ it is still possible to decode time with high accuracy, down to the limiting resolution set by the 100 ms timebin of the analysis, even after the linear component has been removed. Error bars show two standard deviations.

## 5 Low cumulative dimensionality improves generalization

Monkeys performing working memory tasks are able to store task relevant variables across the delay period in a way that generalize to new delay intervals. For example, if the duration of the delay interval is increased the monkey will generalize to the new task without needing to retrain. This generalization ability places constraints on the types of neural dynamics that support working memory. In the data, we observe that neural trajectories with high cumulative dimensionality do not allow for good generalization, suggesting a monkey relying on these dynamics would need to retrain to adjust to a longer delay interval. In contrast, data with low cumulative dimensionality enables computations learned at one point in time to generalize to other points in time.

The cumulative dimensionality over time, after the offset of the external event, is shown in Figure 6 (see supplementary Figure S2 for the RNN models). For all datasets the dimensionality increases much slower than in the case of the stabilized reservoir network^10^, which explores new dimensions of state space at each point in time. After an initial rapid increase, which terminates around 500 ms in all datasets, the cumulative dimensionality increases very slowly or saturates. The initial rapid increase reflects the ability to decode time with high precision, which is probably due to a relatively fast transient that follows the offset of the stimulus. The slow increase observed in the remaining part of the delay is consistent with the strong linear ramping component observed in Figure 5 as ramping activity would cause the neural trajectory to lie along a single line in state space and so the cumulative dimensionality would be one. The cumulative dimensionality is stable as the number of timepoints along the neural trajectory are varied as shown in supplementary Figures S3-S5.

**Figure 6:**
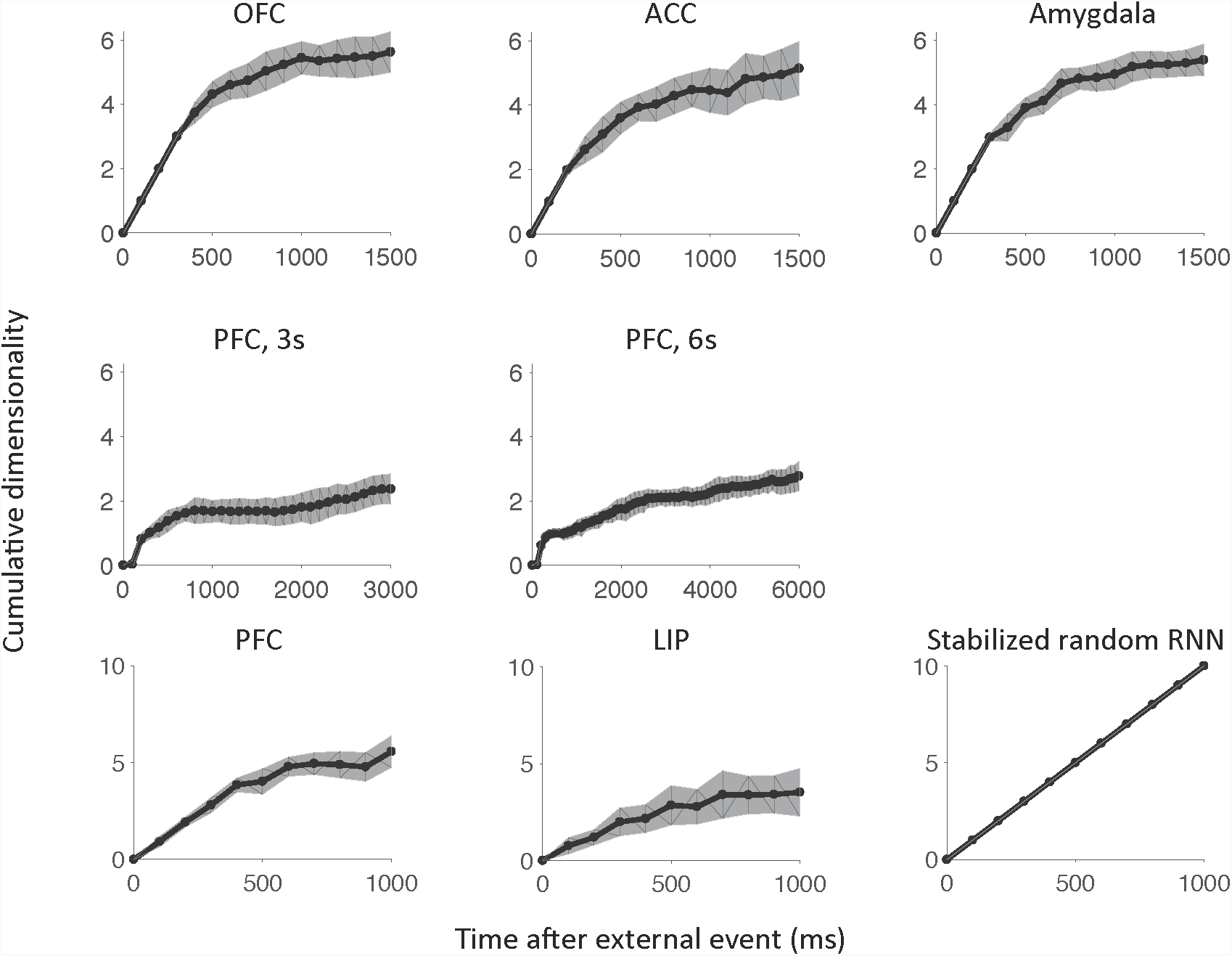
The cumulative dimensionality of the neural activity increases slowly over time after a transient of approximately 500 ms. In contrast, the cumulative dimensionality increases linearly in the stabilized random network^10^. The top two rows show dimensionality for tasks in which timing is not explicitly important. The bottom row shows dimensionality for tasks in which timing is required. Error bars show one standard deviation.

The low dimensional ramping trajectories seen in the neural data may offer computational benefits, allowing computations to generalize across time^31^. For example, consider the task of Saez et al.^15^ where the offset of a visual stimulus is followed by a delay period and then either a water reward or no reward. Imagine a neuron learns to linearly combine neural activity from prefrontal cortex after the offset of the visual stimulus in order to predict whether a reward will be delivered. After some time has elapsed, e.g. a second, will this same neuron still be able to correctly predict the upcoming reward? Will this same linear combination of information from prefrontal cortex still be useful at a different time? The answer depends on how the neural dynamics evolve. The low dimensional ramping trajectories of the neural data allow a linear classifier trained at a few points in time to have predictive power at other points in time (Figure 7). In particular, in OFC, ACC and amygdala (Figure 7, top row), a linear classifier trained to decode the value of the stimulus on a fraction of timepoints can decode the value with an accuracy close to 100% also at the other timepoints. In PFC, a classifier trained to decode high versus low vibro-tactile stimuli can also generalize across time, though the decoding performance is lower than in the case of reward decoding. This is compatible with the stability of the geometry of neural representations observed in Spaak et al.^31^ and with the ability of a linear readout to generalize across experimental conditions observed in Bernardi et al.^16^. This ability to generalize to other timepoints is observed also in the RNN model trained with backpropagation (Figure 7, see all the plots with the label ‘RNN’). In contrast, for the high dimensional neural dynamics of the simulated stabilized random RNN, a linear classifier trained at a few points in time performs near chance level at other points in time (Figure 7, bottom row).

**Figure 7:**
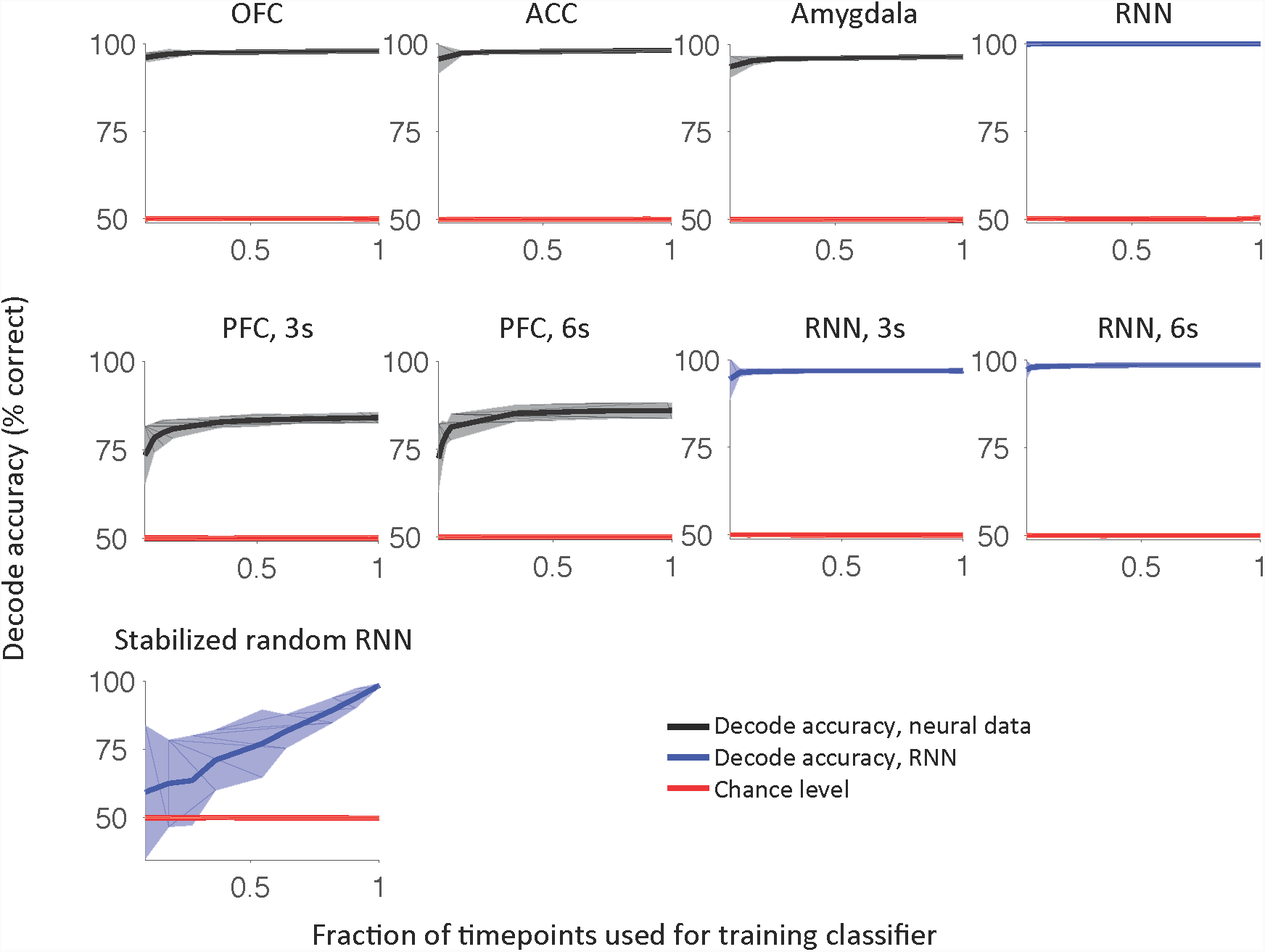
Decode generalization when classifying neural activity with low and high cumulative dimensionality. The decode accuracy of a binary classifier (colored in black for data and blue for RNN models) is shown as the number of timepoints used during training is varied. The chance level is shown in red. The neural activity with low cumulative dimensionality (fixed point dynamics in the top row and ramping activity in the middle row) allows a classifier trained at a single timepoint to perform with high accuracy when tested at other times. This is in contrast to neural activity with high cumulative dimensionality (bottom row) where a decoder trained at a single timepoint performs at chance level when tested at other timepoints. To assess generalization performance the classifier is always tested on timepoints from the entire delay interval with the exception of the first 500 ms after stimulus offset for the neural datasets. In the top row, the decoder classifies rewarded versus non-rewarded trials. In the middle row, the decoder classifies high versus low frequencies. In the bottom row, the decoder classifies trials from the two patterns the network is trained to produce in Laje and Buonomano^10^. The plotted decode accuracy is the mean of the classifier performance across this interval. Error bars show two standard deviations.

## 6 Discussion

Delay activity is widely observed in cortical recordings, and is believed to be important for two functions that could be difficult to combine in the same neural circuit: the first, is to preserve information robustly against the flow of time (working memory); the second, is to actually track the passage of time to anticipate stimuli and plan future actions. To understand the mechanisms underlying delay activity we introduced two main analyses, namely, decoding the passage of time from neural data and computing the cumulative dimensionality of the neural trajectory as it evolves over time. These two analyses allow us to disambiguate different classes of neural dynamics. Our analysis of four datasets revealed that it is possible to decode time, but only with limited precision in tasks where timing information is irrelevant. The precision is significantly higher when timing information is important for performing the task. The dynamics of the neural activity is low dimensional and the ability to decode time relies on the ramping component of the activity. This is true when a transient of approximately 500 ms following the offset of the stimulus is excluded from the analysis. During these 500 ms it is likely that the activity is still driven by the sensory input, and it does not reflect the internal dynamics of the neural circuits. These results indicate that the experimental observations are more compatible with low dimensional dynamical models like the recurrent neural network that we proposed, rather than chaotic dynamics, for which the dimensionality would grow linearly with time. Previous studies^26^ of the vibrotactile stimulation experiment^22^ that we also analyzed show that it is not easy to reproduce the data using chaotic networks similar to those reviewed by Buonomano and Maass^13^. Moreover, recent experiments on rodents ^20, 32^ also show that the trajectories in the firing rate space are low dimensional, with a time varying component that is dominated by ramping activity.

All these results require some discussion. It is possible that the dynamics are chaotic but with an autocorrelation time that is relatively long, comparable with the entire delay interval that we considered. In this case the activity would change slowly, preventing the neural circuit from exploring a large portion of the firing rate space in the limited time of the experiment. The cumulative dimensionality would still grow linearly, but on a much longer timescale, and on the timescale of the experiment, it would be approximately constant. Although possible, this scenario would have to assume that the autocorrelation time can vary on multiple timescales in order to explain the rapid variations observed during the initial transients and, at the same time, the very slow variations observed later, when the cumulative dimensionality stops growing.

One of the robust results of our analysis is that the observed trajectories in the firing rate space are low dimensional. This seems to be in contrast with other studies in which the dimensionality of the neural representations was reported to be high (see e.g. 33–35) or, as high dimensional as it could be^36^. However, it is important to stress that the dimensionality measured in these other studies is a static dimensionality: it is the minimal number of coordinate axes needed to determine the position of all the points in the firing rate space that correspond to different conditions of the experiment (e.g. a single condition in the task of Romo et al.^22^ is all trials with the same vibrotactile frequency). The firing rates of the different conditions are all estimated in the same time bin. In our case, we considered the points that correspond to different time bins for the *same* condition. So it seems that the dimensionality across different conditions is usually high, whereas the trajectories corresponding to each condition is low. This is not surprising given that high dimensionality across conditions is needed in tasks like the one studied in Rigotti et al.^33^, whereas it is probably not required for the delay activity trajectories that we analyzed here or in other situations in which the task relevant variables do not need to be mixed non-linearly (see e.g. 16). To maximize the ability to generalize, the dimensionality should always be the minimal required by the task, and this is probably the case also in the tasks that we analyzed.

Reservoir networks are constructed to perform difficult tasks in which time and many other quantities (e.g. combinations of events occurring at different times) can be decoded using a simple linear decoder. In the tasks that we considered, there is probably no need for such high dimensional trajectories. The observation that the recurrent neural network (RNN) models that we trained with backpropagation generate low dimensional trajectories is an indication that high dimensionality is not needed. And low dimensional trajectories allow for better generalization as we showed in Figure 7. It is important to note that in reservoir networks time can be decoded with a simple linear readout. This is probably not the case for the low dimensional trajectories that we observed (indeed, our time decoder illustrated in Figure M2A is non-linear). However, there are situations in which a non-linear decoder is not required to be able to decode time in certain time bins. For example, even in the low dimensional case in which a trajectory is perfectly linear, it is often possible to linearly separate the last point from the others. So a linear readout would be able to report that a certain time bin is at the end of a given interval, and “anticipate” the arrival of the second stimulus. For more complex computations on the low dimensional trajectories, the brain might employ a non-linear decoder, which could easily be implemented by a downstream neural circuit that involves at least one hidden layer.

The RNN models that we trained using backpropagation through time (BPTT) reproduce many of the important features of the four datasets that we analyzed. RNNs have also been successfully used to model the ready-set-go task when analyzing the motor production interval between the set and go cues^37, 38^, whereas in this work we analyze the interval when time is initially encoded, between the ready and set cues. The success of these simulated RNN models is surprising given that BPTT is an artificial algorithm, for which there is no comparable biologically plausible implementation at the moment. Methods have been proposed for biologically plausible learning in recurrent networks^39, 40^ but it should not be assumed that these methods scale to harder problems, as this scaling has proven difficult for biological approximations to backpropagation for feedforward networks^41^. However, the brain and the simulated recurrent neural networks that we built share similar constraints as they are both trained to perform the same tasks efficiently in the presence of noise. This is probably why some of the features of the neural representations are similar. It remains possible that some of the important mechanisms are actually implemented in a very different way. For example, the ramping activity might be a consequence of some biochemical processes that are present at the level of individual neurons or synapses in the biological brain^42, 43^, but not explicitly modelled in the recurrent neural network, in which all the elements are simple rate neurons. This process can be imitated in the network by tuning the weights between neurons of canonical circuits that essentially are devoted to implementing a specific biochemical process. A more complex analysis will be developed to reveal these canonical circuits. In the meantime it is important to keep in mind that we do not necessarily expect a one-to-one correspondence between the neurons in the RNN and the neurons in the brain.

A fundamental challenge in studying neural activity that evolves over time is understanding *what* computational capabilities can be supported by the activity and *when* these dynamics change to support different computational demands. Our time decode and cumulative dimensionality analyses offer a tool for parcellating neural activity into computationally distinct regimes across time by objective classification of electrophysiological activity. In this work we apply these analyses to delay period activity and find that low dimensional trajectories provide a mechanism for the brain to solve the problem of time invariant generalization while retaining the timing information necessary for anticipating events and coordinating behavior in a dynamic environment.

## Acknowledgments

We thank members of the Center for Theoretical Neuroscience at Columbia University for useful discussions. Research supported by NSF NeuroNex Award DBI-1707398, NIH training grant 5T32NS064929 (CJC), the Simons Foundation, the Gatsby Charitable Foundation and the Schwartz Foundation. RR was supported by the Dirección General de Asuntos del Personal Académico de la Universidad Nacional Autónoma de México (PAPIIT-IN210819) and Consejo Nacional de Ciencia y Tecnologίa (CONACYT-240892). AG was supported by the Division of Intramural Research of the NIMH (Z01MH-01092) and by the italian FIRB 2010 grant (RBFR10G5W9 001). AG wishes to thank Steve Wise, Satoshi Tsujimoto and Andrew Mitz for their numerous contributions.

## Methods

### Decoding time

To create the “two-interval time decode matrix”, as detailed in Figure M1, we first subdivide the time after stimulus offset into nonoverlapping 100 ms intervals. We take the vector of firing rates recorded from all neurons during a single interval (interval *i*) and train a logistic regression classifier to discriminate between this and another interval (interval *j*). We test the classifier on held-out trials and record the performance. This number, between 50% and 100%, from the binary classifier trained to discriminate intervals *i* and *j* is recorded in pixel (*i,j*) of the “two-interval time decode matrix.” If the decode accuracy is 100% the pixel is colored yellow and if the decode accuracy is 50% the pixel is colored blue. We use 3/5 of the trials for training and the remainder for testing, resampling single unit recordings to create 10,000 trials for training and testing and then performing cross-validation 100 times to establish the final mean decode accuracy.

**Figure M1:**
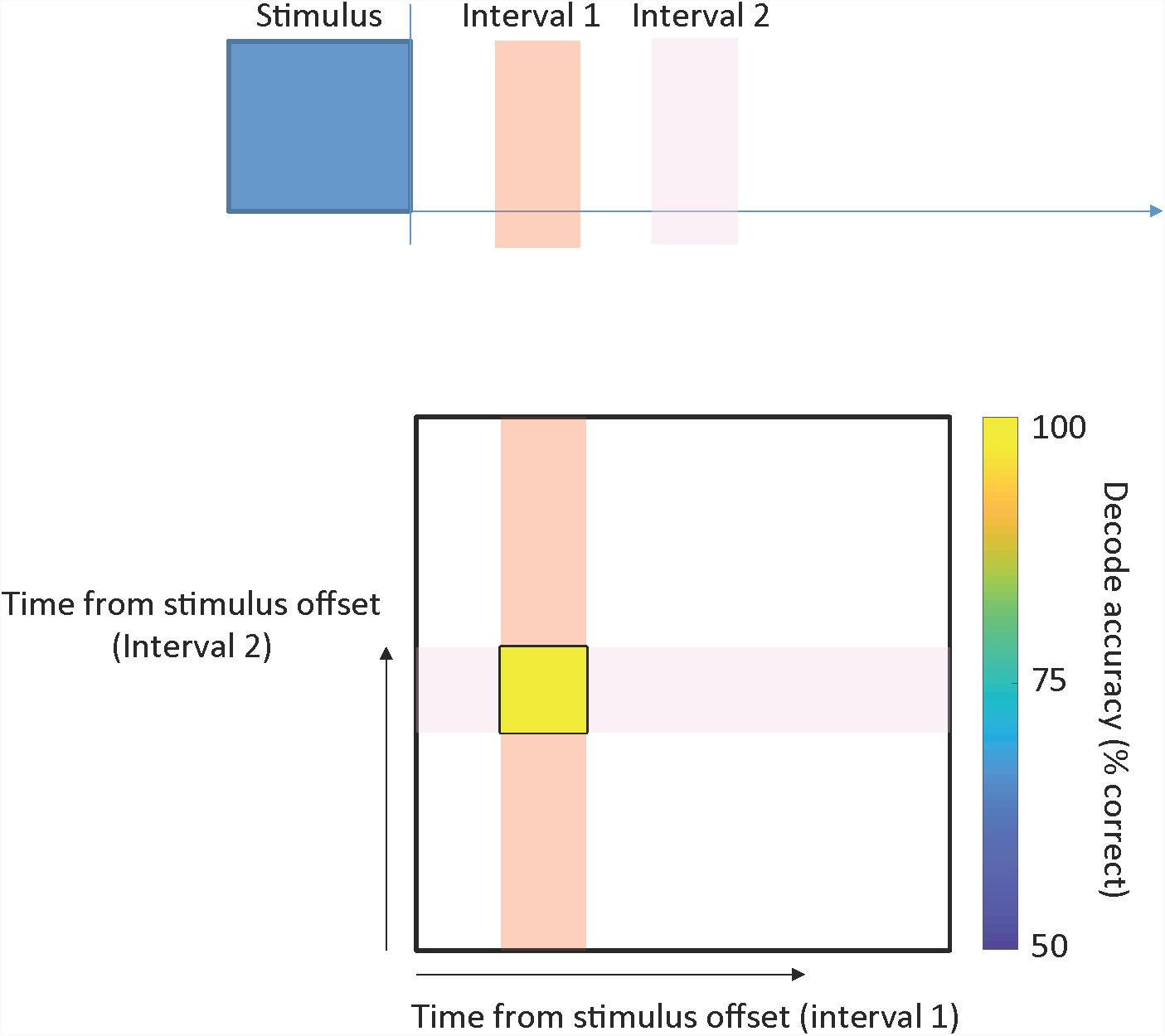
Two-interval time decode matrix. Subdivide the time after stimulus offset into nonoverlapping intervals. Take the vector of firing rates recorded from all neurons during a single interval (interval 1, for example) and train a binary classifier to discriminate between this and another interval (interval 2). Test the classifier on held-out trials and record the performance. This number, between 50% and 100%, from the classifier trained to discriminate intervals *i* and *j* is recorded in pixel (*i,j*) of the “two-interval time decode matrix.” If the decode accuracy is 100% the pixel is colored yellow (as shown in this example) and if the decode accuracy is 50% the pixel is colored blue.

To quantify the timing uncertainty of the neural data at a given point in time we train a classifier to predict the time this recording was made (Figure M2A), and then use the spread of predictions when classifying firing rates from different trials as our measure of timing uncertainty. This multi-class classifier takes the firing rates from all neurons at a given timepoint (time is discretized in 100 ms bins) and predicts the time this recording was made. This is in contrast to the two-interval time decode analysis where the binary classifier only discriminates between two timepoints; the multi-class classifier attempts to predict the actual timepoint within the trial, e.g. 1000 ms after stimulus offset. We classify 10,000 trials (obtained through resampling single unit recordings) at each point in time yielding a distribution of predictions around the true value (Figure M2B). We calculate the standard deviation of this distribution (Figure M2C) and this is the metric for timing uncertainty. To classify trials we combine the pairwise binary classifications from the “two-interval time decode matrix” (Figure M2A), however, we obtain similar predictions with other multi-class classifiers. The chance level for the timing uncertainty is computed by training and testing the classifier on neural data with random time labels. To gain some intuition about the chance level distribution we can imagine the classifier predictions, *X*, are random and uniformly distributed over some interval *T*. We can compute various properties of *X*, for example, the expectation of *X* is *E*[*X*] = 0.5 *\** (min(*T*) + max(*T*)). Its standard deviation is 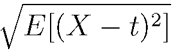. If the chance-level-classifier guesses uniformly on the interval *T* then *E*[(*X − t*)^2^] = *t*^2^ − *t \** (min(*T*)+ max(*T*))+ ((max(*T*)^3^) - min(*T*)^3^)*/*(3 *\** (max(*T*) - min(*T*))), yielding a U-shaped curve for the timing uncertainty, 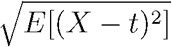, as seen in the chance level curves of Figures 3-5. Intuitively, the chance level is U-shaped as a classifier with uniform, random predictions can make larger errors when the true value is at the edge of the interval. In practice, we don’t use the analytic expression for 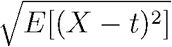 but calculate this after training and testing a classifier on neural data that has the timepoint labels randomly shuffled. We use 3/5 of the trials for training and the remainder for testing, resampling single unit recordings to create 10,000 trials for training and testing the classifier and then performing cross-validation 100 times.

### Neural dimensionality

Our measure of neural dimensionality quantifies the *stable* component of the neural trajectory across trials (Figure M3). The stable component of the neural trajectory is important as it can be used for consistent computations by downstream neurons. Other common measures of dimensionality yield a large dimensionality even for random Gaussian noise data, with no consistent firing rate fluctuations across trials, as shown in Figure M4. For this data, the only consistent aspect is the mean and our ‘trajectory reconstruction dimensionality’ quantifies this data as zero dimensional.

To compute the cumulative neural dimensionality over time we used a cross-validation procedure to estimate the number of dimensions that gave us the greatest predictive power on firing rates from held out data. We first subdivided the time after stimulus offset into *T* nonoverlapping 100 ms intervals. For each timepoint, from 1 through *t* = 1,…, *T* that we wished to estimate the cumulative dimensionality we constructed a training and test matrix (FRtrain and FRtest) of size number-of-neurons *× t* containing firing rates after averaging spikes in 100 ms bins and across trials. FRtrain and FRtest contained averages from nonoverlapping sets of trials. If the neural activity was the same on every trial then FRtrain and FRtest would be equal and we would be able to predict the firing rates in FRtest perfectly from FRtrain. However, there is variability that is not shared between FRtrain and FRtest so FRtrain is not perfectly predictive. We also expect some variability in the neural trajectory of FRtrain is shared with FRtest so there is an optimal subspace of FRtrain that will yield the greatest prediction accuracy for FRtest. We estimated this subspace by first projecting FRtrain onto principal components 1 through *k*, sorted in order of descending variance so principal component 1 captures the most variance. We define the dimensionality as the number of principal components *k* that yields the greatest predictive accuracy for FRtest, i.e. that yields the smallest squared error between FRtest and the ‘denoised’ trajectory of FRtrain after projecting onto the first *k* principal components (Figure M3). We used 3/5 of the trials for training and the remainder for testing, repeating cross-validation 200 times. Single unit recordings were resampled to create 1000 trials for the FRtrain and FRtest matrices.

**Figure M2:**
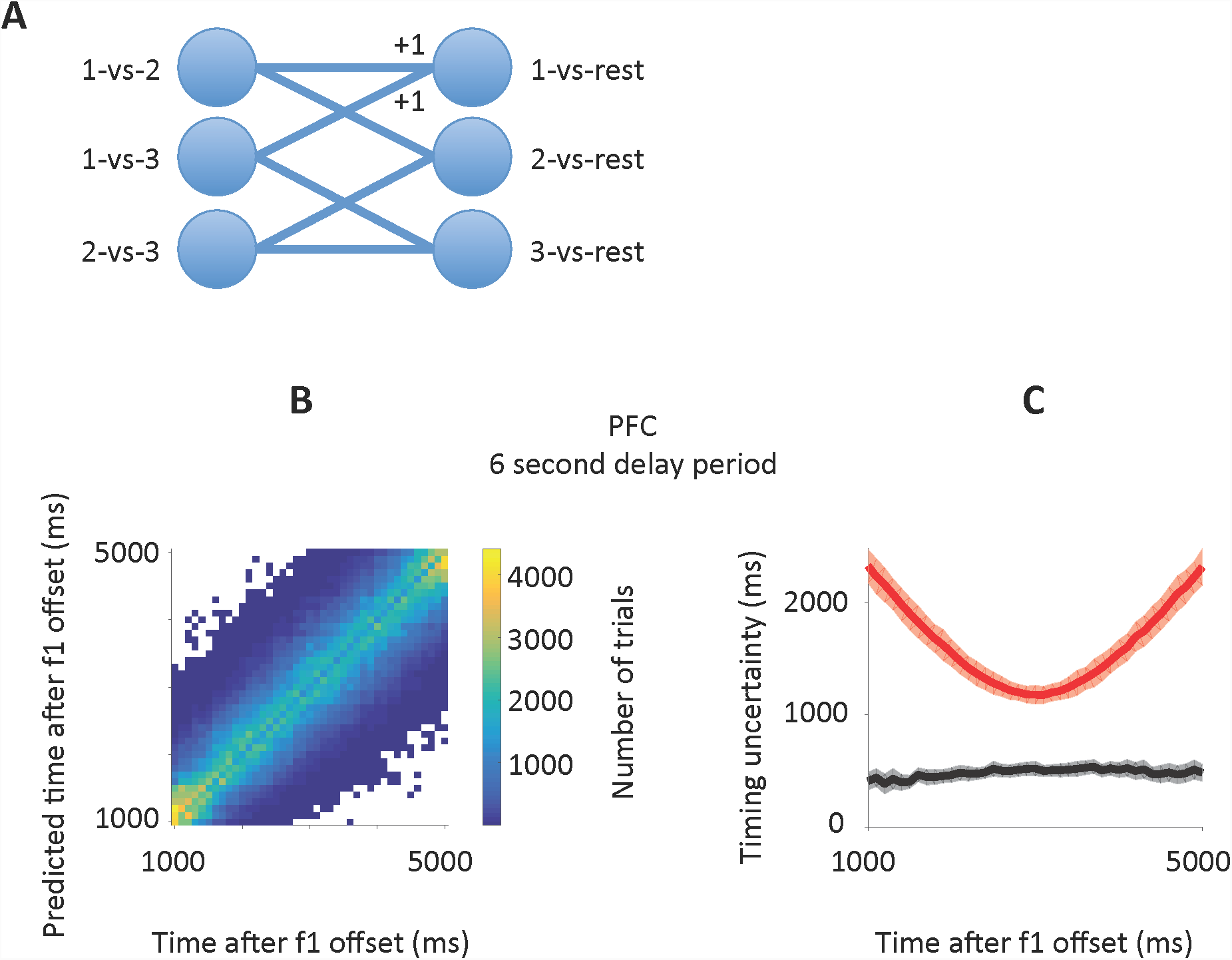
Computing the timing uncertainty. **(A)** To classify neural data into one of *T* timebins we linearly combine results from the pairwise binary classifications obtained from the two-interval time decode. A schematic for *T* = 3 is shown here. For each datapoint, i-vs-j in [0,1] is the confidence of the binary classifier discriminating classes i and j in favor of the former class. The confidence of the classifier for the latter class is computed by j-vs-i = 1 i-vs-j. To obtain the final classifier prediction we add the confidence votes. For example, the total confidence in favor of timepoint 1 (1-vs-rest) is the sum of 1-vs-2 and 1-vs-3. A similar calculation is performed for the total confidence in favor of the other timepoints (2-vs-rest and 3vs-rest), and the timepoint with highest confidence is the output of the multi-class classifier. **(B)** To quantify the timing uncertainty at each point in time we calculate the predicted time (y-axis) relative to the actual elapsed time from stimulus offset (x-axis). The predicted time (cross-validated) is calculated for each of the 10,000 trials (obtained through resampling single unit recordings) and the results are shown as a heatmap. Data is from the delay interval of Romo et al.^22^ **(C)** The standard deviation of the distribution in (B) is shown in black and is our metric for timing uncertainty. The chance level is shown in red. Error bars show two standard deviations.

**Figure M3:**
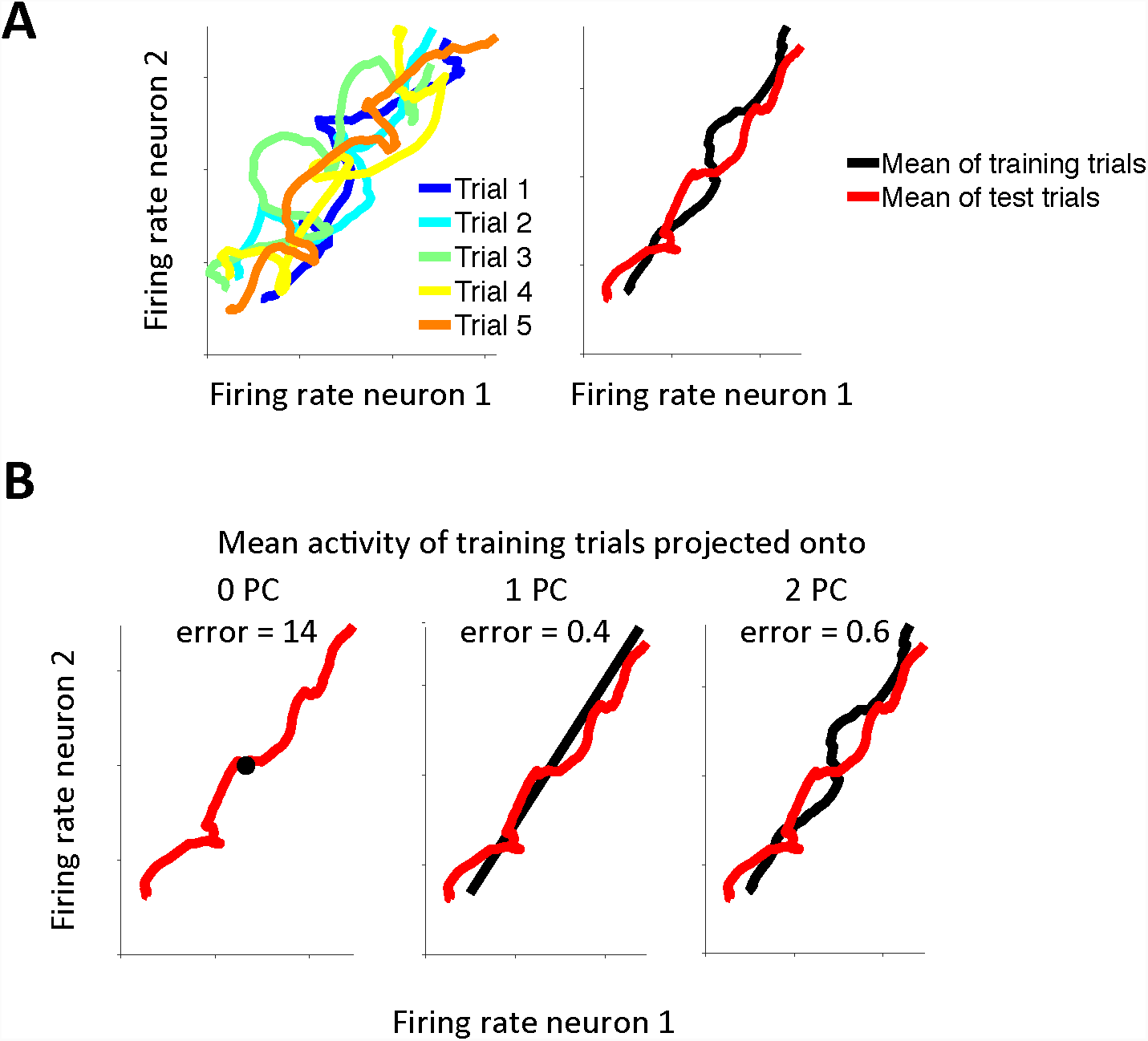
Computing cumulative dimensionality. **(A)** The figure on the left shows five trials recorded from an example dataset consisting of two neurons. These trials are split into training and test groups, and then averaged within groups, as shown in the figure on the right. **(B)** Principal components are calculated from the mean training trajectory. The mean training trajectory is then projected onto the mean activity (left), first principal component (center), up to the maximum number, which in this example is the first two principal components (right). The mean squared error is then calculated between the training trajectory, after the projections, and the test trajectory for all timepoints. In this example, the projection of the training trials onto only one principal component yields the lowest error, i.e. is best able to predict the test trajectory, and so the dimensionality is one. This process is repeated for neural trajectories of varying lengths in order to assess the dimensionality as the firing activity evolves over time.

**Figure M4:**
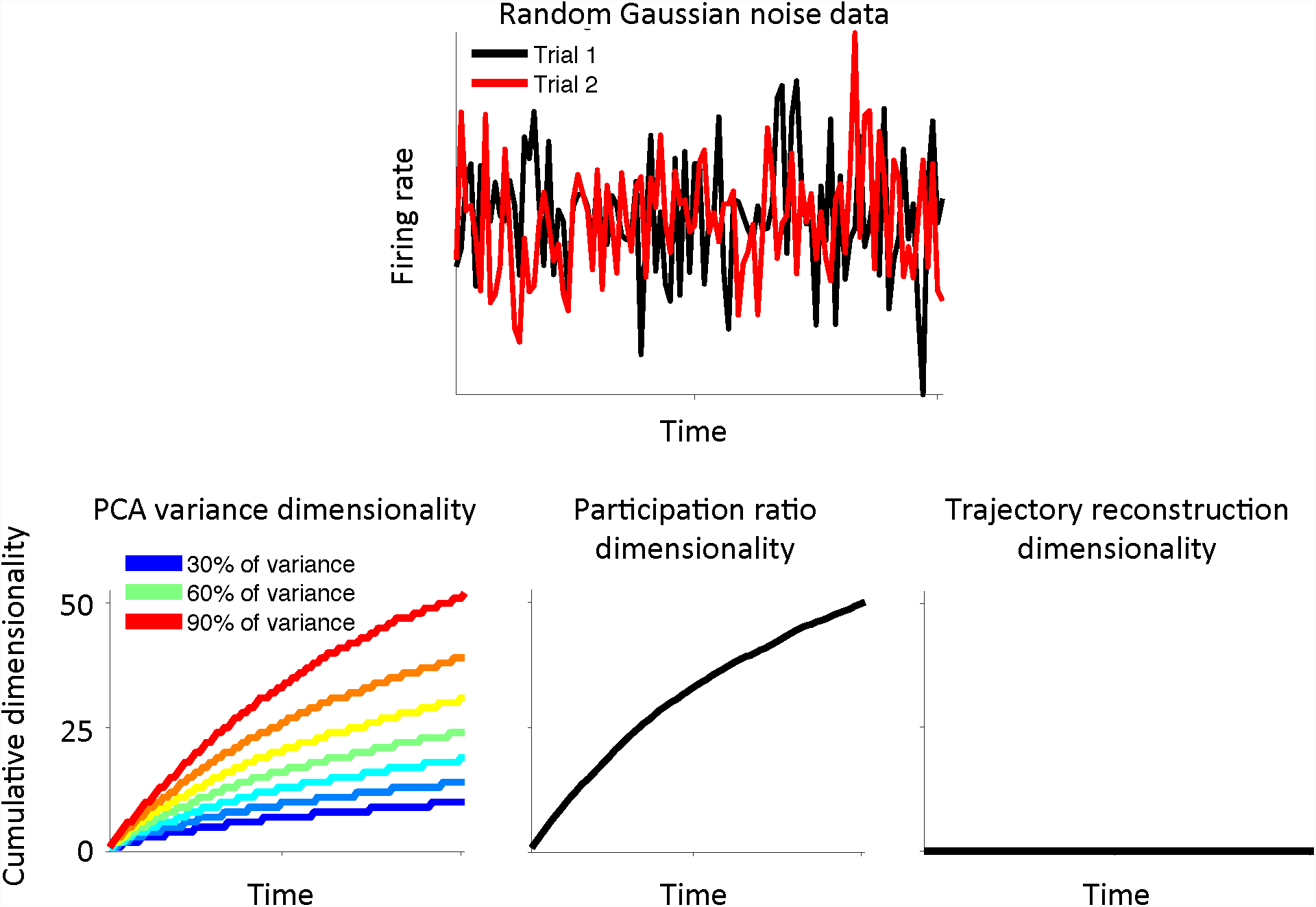
Cumulative dimensionality of random Gaussian noise data. The cumulative dimensionality is the dimensionality required to exaplain timepoints 1 through *t* where *t* = 1, 2, 3, *…* is increased from 1 to some maximum value. In the figure on the left, the dimensionality is quantified as the number of principle components required to explain a fraction of the variance (e.g. 90% as shown in red). The center figure shows the cumulative dimensionality when using the Participation Ratio^44, 45^. Both methods show an increase in the dimensionality over time. However, by construction, there are no stable trajectories across trials that can be used for consistent computation. The only consistent aspect of the data is the mean and our ‘trajectory reconstruction dimensionality’ quantifies this data as zero dimensional.

### Decode generalization

We first subdivide the delay period into *T* nonoverlapping 100 ms intervals, excluding the first few hundred milliseconds after stimulus offset to better study the intrinsic dynamics of the delay period without transient activity caused by the offset of the visual stimulus. To quantify the ability of a classifier to generalize to other timepoints we train a logistic regression classifier on a fraction of timepoints (from near 0 to 1) and then test its accuracy across the entire interval (see Figure M5). The plotted decode accuracy in Figure 7 is the mean of the classifier performance across the entire interval. In the top row of Figure 7, the decoder classifies rewarded versus non-rewarded trials in the dataset of Saez et al.^15^ during the interval from 500 ms to 1500 ms after stimulus offset. In the middle row, the decoder classifies high versus low frequency trials in the dataset of Romo et al.^22^ during the interval from 500 ms to 3000 ms after stimulus offset (3s delay period) and 500 ms to 6000 ms (6s delay period). In the bottom row, the decoder classifies trials from the two patterns the network is trained to produce in Laje and Buonomano^10^ during the interval from 200 ms to 1200 ms after stimulus offset. The chance level is computed by randomly shuffling these labels before training and testing the classifier. We use 3/5 of the trials for training and the remainder for testing, resampling single unit recordings to create 10,000 trials for training and testing the classifier and then performing cross-validation *T* times. Each time cross-validation is performed a new, unique set of timepoints are randomly chosen and used for training the classifier. Note that when only a single timepoint is used for training the classifier, we cycle once through each and every timepoint in the interval. Error bars show two standard deviations.

**Figure M5:**
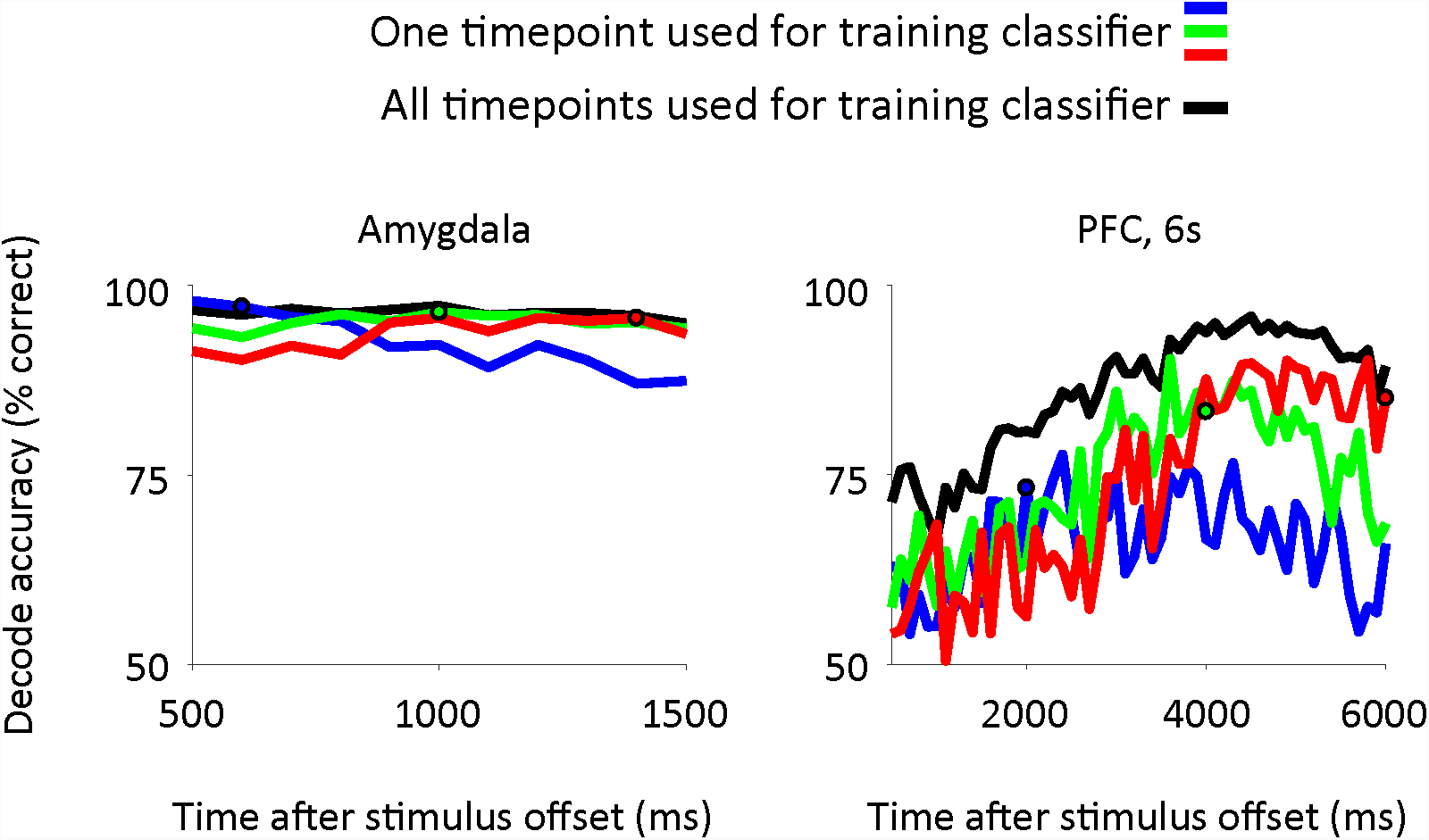
Decode accuracy when training the classifier on one timepoint (blue, green, and red curves) and all timepoints (black curve). The decode accuracy is shown for a binary classifier trained to discriminate neural data from rewarded versus non-rewarded trials for the task of Saez et al.^15^ (Amygdala, left) and high versus low frequency trials during the task of Romo et al.^22^ (PFC, 6s on right). Three examples are shown when the classifier is trained at a single timepoint (highlighted with circles); the classifier is trained at times 600 ms (blue), 1000 ms (green) and 1400 ms (red) after stimulus offset for the Amygdala and at times 2000 ms (blue), 4000 ms (green), and 6000 ms (red) after stimulus offset for the PFC. The decode accuracy is computed using trials that were not used for training the classifier. The leftmost datapoint in Figure 7, for example, summarizes the decode generalization when the smallest fraction of timepoints are used for training the classifier and is computed as the mean decode accuracy of all the curves trained at a single timepoint (as shown), along with all other curves obtained during the other iterations of cross-validation (not shown).

### Model description

For each of the four working memory tasks we train a recurrent neural network (RNN) model. Our network models consist of a set of recurrently connected units (*N* = 100). The dynamics of each unit in the network *u*_*i*_(*t*) is governed by the standard continuous-time RNN equation:

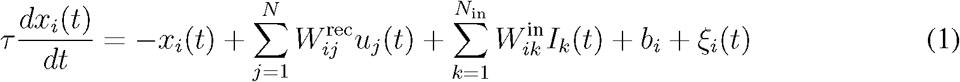

for *i* = 1, *…, N*. The activity of each unit, *u*_*i*_(*t*), is related to the activation of that unit, *x*_*i*_(*t*), through a nonlinearity which in this study we take to be *u*_*i*_(*t*) = tanh(*x*_*i*_(*t*)). Each unit receives input from other units through the recurrent weight matrix *W* ^rec^ and also receives external input, *I*(*t*), that enters the network through the weight matrix *W* ^in^. Each unit has two sources of bias, *b*_*i*_ which is learned and *ξ*_*i*_(*t*) which represents noise intrinsic to the network and is taken to be Gaussian with zero mean and constant variance. The network was simulated using the Euler method for *T* time steps, with a step size of duration *τ/*10 = 10 ms. To perform tasks with the RNN we linearly combine the firing rates of units in the network and use this as the output. The linear readout neurons, *y*_*j*_(*t*), are given by the following equation:

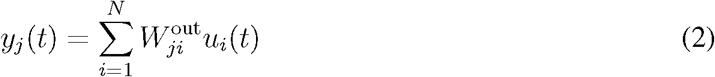

The RNN for each working memory task has the same architecture but the network parameters *W* ^rec^, *W* ^in^, *b* and *W* ^out^ are different for each task and adjusted to accomplish the task-specific transformation of time-varying inputs to time-varying outputs.

We optimized the network parameters *W* ^rec^, *W* ^in^, *b* and *W* ^out^ to minimize the squared error in equation (3) between target outputs and the network outputs generated according to equation (2).

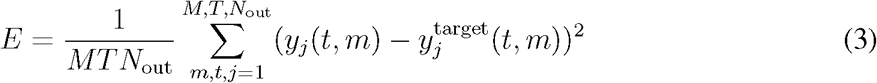

Parameters were updated with the Hessian-free algorithm ^21^ using minibatches of size 500, i.e. 500 sequences of length *T* for each parameter update. In addition to minimizing the error function in equation (3) we regularized the input and output weights according to equation (4).

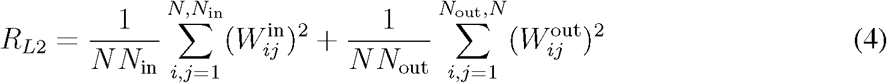

The parameters *W* ^out^ and *b* were initialized to zero. *W* ^in^ was initialized with random values drawn from a normal distribution with zero mean and variance 1*/N*_in_. *W* ^rec^ was initialized as a random orthogonal matrix ^46^.

For all four networks, the “firing rate” of each of the 100 units (*u*(*t*)) is stored every 100 ms and this activity is used for all subsequent analyses.

### Inputs and outputs for the network

The RNN for the context dependent working memory task of Saez et al.^15^ has four inputs (stimulus A, stimulus B, reward, and the end-of-trial cue) and two outputs (reward predictive output and a no-reward predictive output) as shown in Figure M6. During context 1 stimulus A is followed by a reward and stimulus B is not rewarded. During context 2 the associations are reversed and stimulus B is rewarded while stimulus A is not rewarded. Context is not given to the RNN but must be inferred from the previous stimulus/reward pairing stored in the network’s firing activity. The RNN indicates its knowledge of context by switching the appropriate output from zero to one, after stimulus offset, to indicate either an expected future reward or no reward. The inputs are presented serially, e.g. stimulus A, followed by the reward, followed by the end-of-trial cue, each having a value of one for 200 ms before returning to their baseline values of zero. The reward predictive output turns on immediately after the presentation of the stimuli the network thinks will be rewarded and stays on until the end-of-trial cue. The no-reward predictive output turns on immediately after the stimuli the network thinks will not be rewarded and stays on until the end-of-trial cue. The reward can either follow stimulus A (context 1) or stimulus B (context 2). We trained the network using sequences of length 7000 ms with randomly switching contexts and intervals between events. An example trial is shown in Figure M7. During training, the interval between the end-of-trial cue and new stimulus was uniformly distributed between 0 and 1000 ms. The interval between the end of a stimulus and the reward, if present, was uniformly distributed between 0 and 1500 ms. The interval between the end of the reward and end-of-trial cue was uniformly distributed between 0 and 500 ms.

**Figure M6:**
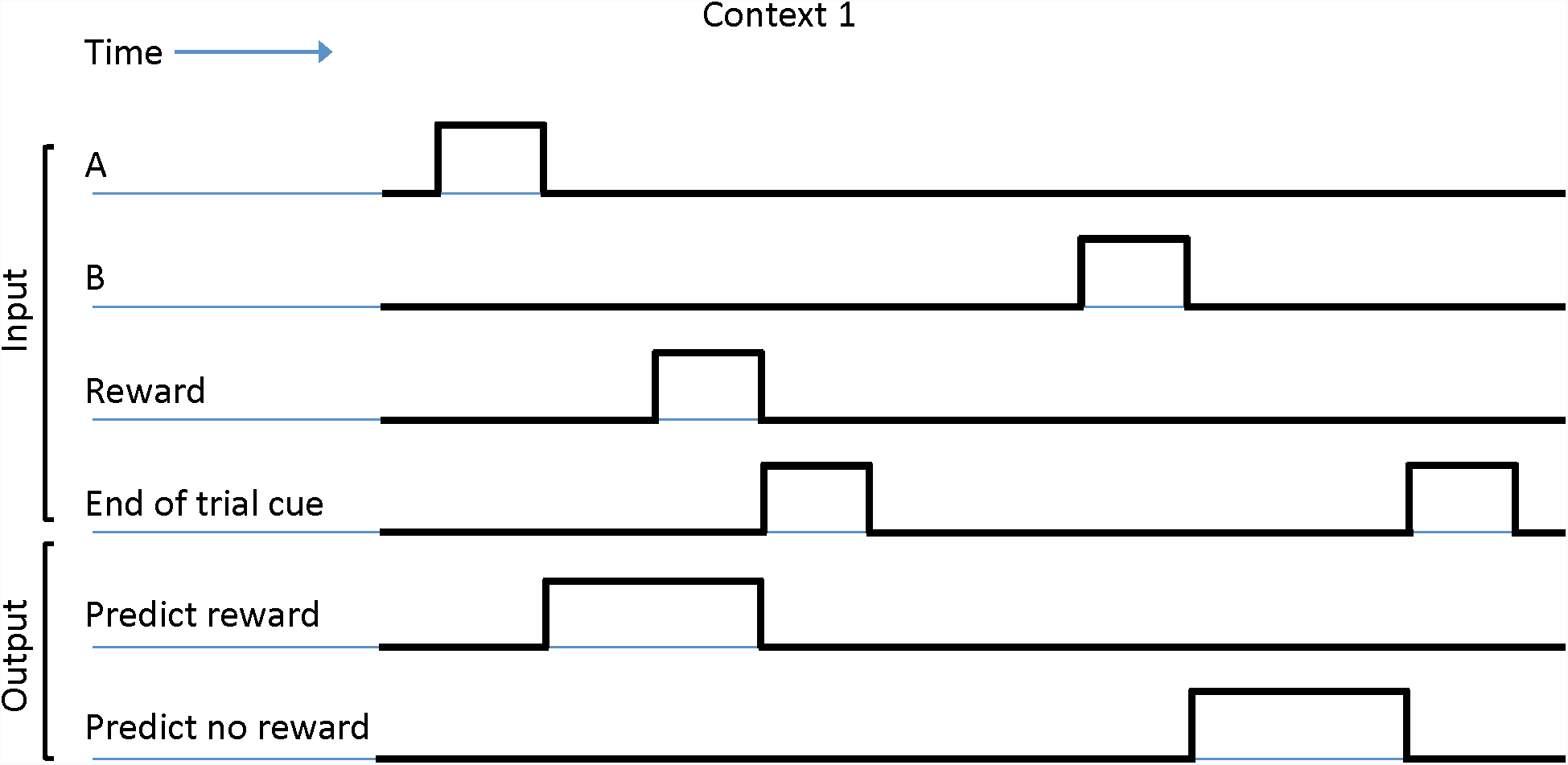
RNN inputs and outputs for the context dependent working memory task of Saez et al.^15^. The RNN has four inputs (stimulus A, stimulus B, reward, and the end-of-trial cue) and two outputs (a reward predictive output and a no-reward predictive output). The mapping between stimulus and reward changes depending on context. During context 1 stimulus A is followed by a reward and stimulus B is not rewarded. During context 2 the associations are reversed and stimulus B is rewarded while stimulus A is not rewarded. The RNN’s task is to predict the upcoming reward following the presentation of a stimulus by selecting the appropriate output. Outputs are shown for context 1.

The RNN for the vibrotactile discrimination task of Romo et al.^22^ is trained to report whether the frequency of the second stimulus (f2) is higher or lower than the frequency of the first (f1), and also to anticipate the time when f2 is presented. The RNN has one input that varies in magnitude to represent the frequency of f1 and f2, and two outputs; an output to indicate the choice for the binary discrimination, and an anticipatory-timer output that turns on before f2 onset (Figure M8). The inputs have a duration of 500 ms with amplitudes similar to Barak et al. ^26^ that linearly map the frequencies between 10-34 Hz to inputs between 0.2 and 1.8: input = 0.2 + (1.8 − 0.2) * (*f*−10)*/*(34 − 10) where *f* is the frequency in Hz. The RNN was trained on sequences of duration 15000 ms with successive presentations of f1 and f2 randomly chosen between 10 and 34 Hz. On each sequence the delay between f1 and f2 was fixed, selected uniformly between 200 and 7000 ms, while the intertrial interval between f2 offset and f1 onset was uniformly distributed between 200 and 1000 ms. The RNN output that performs frequency discrimination takes values of +1 if f2 *>* f1 and −1 if f2 *<* f1. This output is zero until f2 onset, whereupon it takes the appropriate nonzero value until f2 offset, at which time the amplitude returns to zero. The anticipatory-timer output is not constrained during the first f1/f2 pairing in a sequence, because the interval between f1 and f2 has not been established for this sequence. During subsequent inputs of f1/f2 the anticipatory-timer output takes a value of one 200 ms before f2 onset and then returns to zero immediately before f2 onset.

The RNN for the ready-set-go interval reproduction task from Jazayeri and Shadlen^29^ has two inputs (ready and set cues) and one output to indicate the interval between ready and set cues as shown in Figure M9. During training, the ready cue is followed by the set cue with an interval selected from a uniform random distribution between 200 and 1100 ms. The RNN output follows the set cue after a delay that matches the elapsed time between the ready and set cues. All inputs and outputs take the value of zero when they are “off” and one for a duration of 110 ms when they are “on”. The RNN was trained on sequences of duration 4500 ms with multiple presentations of the ready and set cues in each sequence.

The goal of the duration-discrimination task from Genovesio et al.^30, 47, 48^ is to compare the duration of two stimuli (S1 and S2) and select the stimulus that was presented for the longest duration. The RNN for this task has four inputs (S1, S2, go-cue for S1/S2 on left/right of screen, go-cue for S1/S2 on right/left of screen) and two outputs to indicate a hand response to either the right or left (Figure M10). The RNN was trained on sequences of length 5000 ms with a single presentation of S1, S2, and go-cue, per sequence. The order of events within a sequence is pre-stimulus period (uniformly distributed between 100 and 500 ms), S1 (uniformly distributed between 100 and 1500 ms), delay period (uniformly distributed between 0 and 1000 ms), S2 (uniformly distributed between 100 and 1500 ms), delay period (uniformly distributed between 0 and 1000 ms), and a go-cue that initiated the RNN output and remained on until until the end of the sequence. In the experiment, the go-cue was the presentation of the two stimuli (S1 and S2) simultaneously on the right and left side of the screen. On each trial the left and right assignment of S1 and S2 was random so the motor response could not be prepared in advance of this go-cue. To mimic this in the RNN we use two go-cues to indicate whether the position of S1 is on the right or left of the screen. The two RNN outputs then correspond to a hand response to either the right or left. The outputs are zero until the time of the go-cue, when the appropriate output becomes one until the end of the sequence.

**Figure M7:**
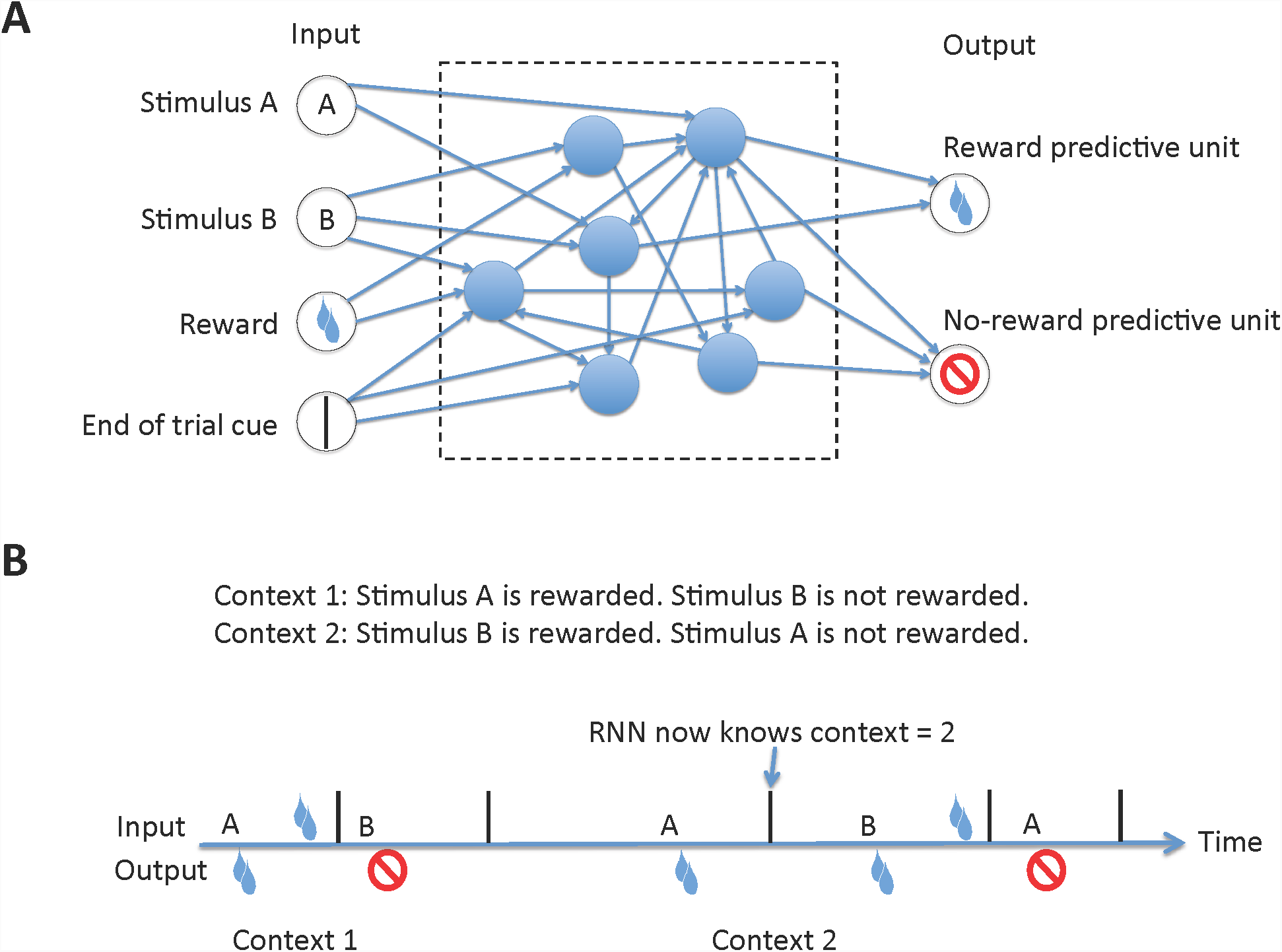
RNN inputs and outputs for the context dependent working memory task of Saez et al.^15^ when the context changes. **(A)** The RNN has four inputs (stimulus A, stimulus B, reward, and the end-of-trial cue) and two outputs (a reward predictive output and a no-reward predictive output). The mapping between stimulus and reward changes depending on context. During context 1 stimulus A is followed by a reward and stimulus B is not rewarded. During context 2 the associations are reversed and stimulus B is rewarded while stimulus A is not rewarded. Note that a single pairing of stimulus/reward or stimulus/end-of-trial-cue is sufficient to determine the context. The RNN’s task is to predict the upcoming reward following the presentation of a stimulus by selecting the appropriate output. **(B)** The inputs and outputs are shown for a single sequence with a change in context. Imagine a preceding stimulus/reward pairing (not shown) has established the context to be 1. Stimulus A and B are inputted and the RNN produces the correct outputs. The context now switches to 2. No explicit contextual cues are given to the RNN so when stimulus A is presented the RNN still responds with the appropriate output for context 1, by activating the reward predictive unit. No reward is inputted, as would be appropriate for context 1, so when the end-of-trial-cue appears the RNN now knows the context has changed to 2. For subsequent inputs the RNN now produces outputs appropriate for context 2.

**Figure M8:**
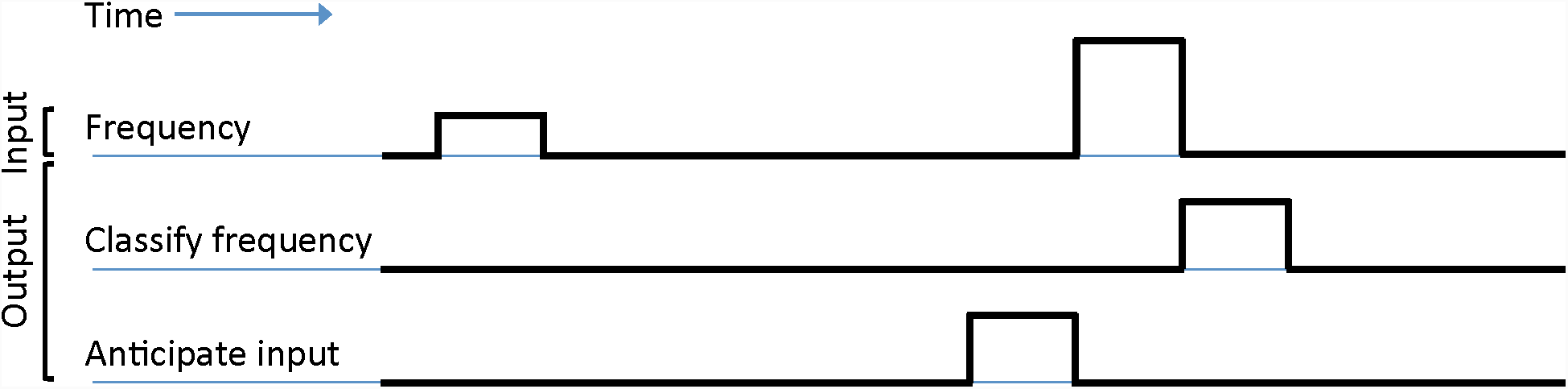
RNN inputs and outputs for the vibrotactile discrimination task of Romo et al.^22^. The RNN has one input representing vibrotactile frequency with the amplitude of an input pulse. After two successive frequency inputs the RNN reports whether the frequency of the second stimulus is higher or lower than that of the first by modulating an output to be +1 or −1 respectively. The RNN also anticipates the timing of the second frequency input by activating a second output before the onset of the stimulus.

**Figure M9:**
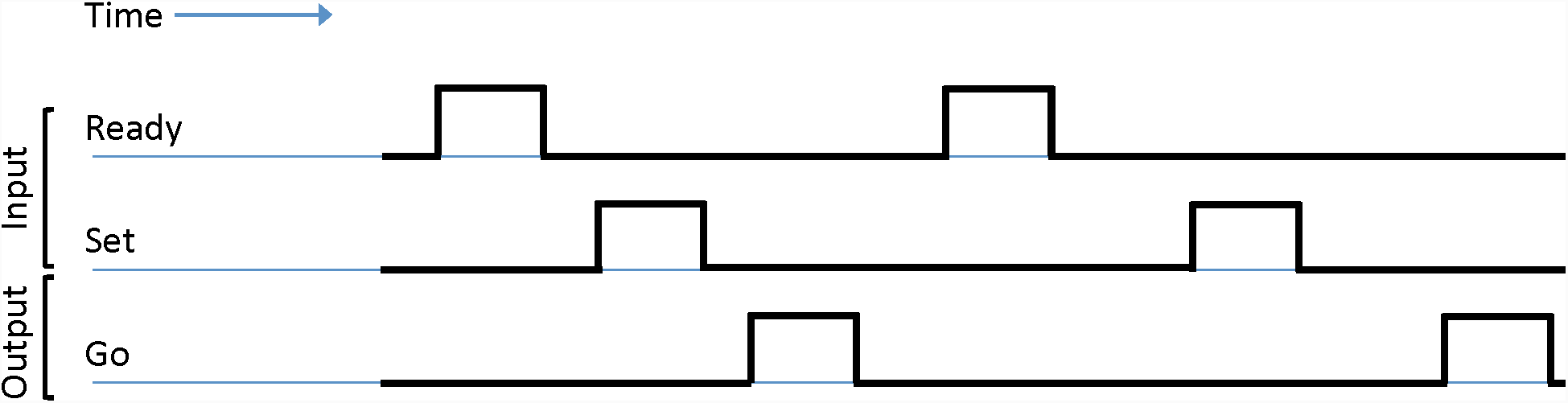
RNN inputs and outputs for the ready-set-go interval reproduction task from Jazayeri and Shadlen^29^. The RNN tracks the duration of the interval between ready and set cues (demarcated by two input pulses) in order to reproduce the same interval with a self initiated output at the appropriate time after the set cue.

**Figure M10:**
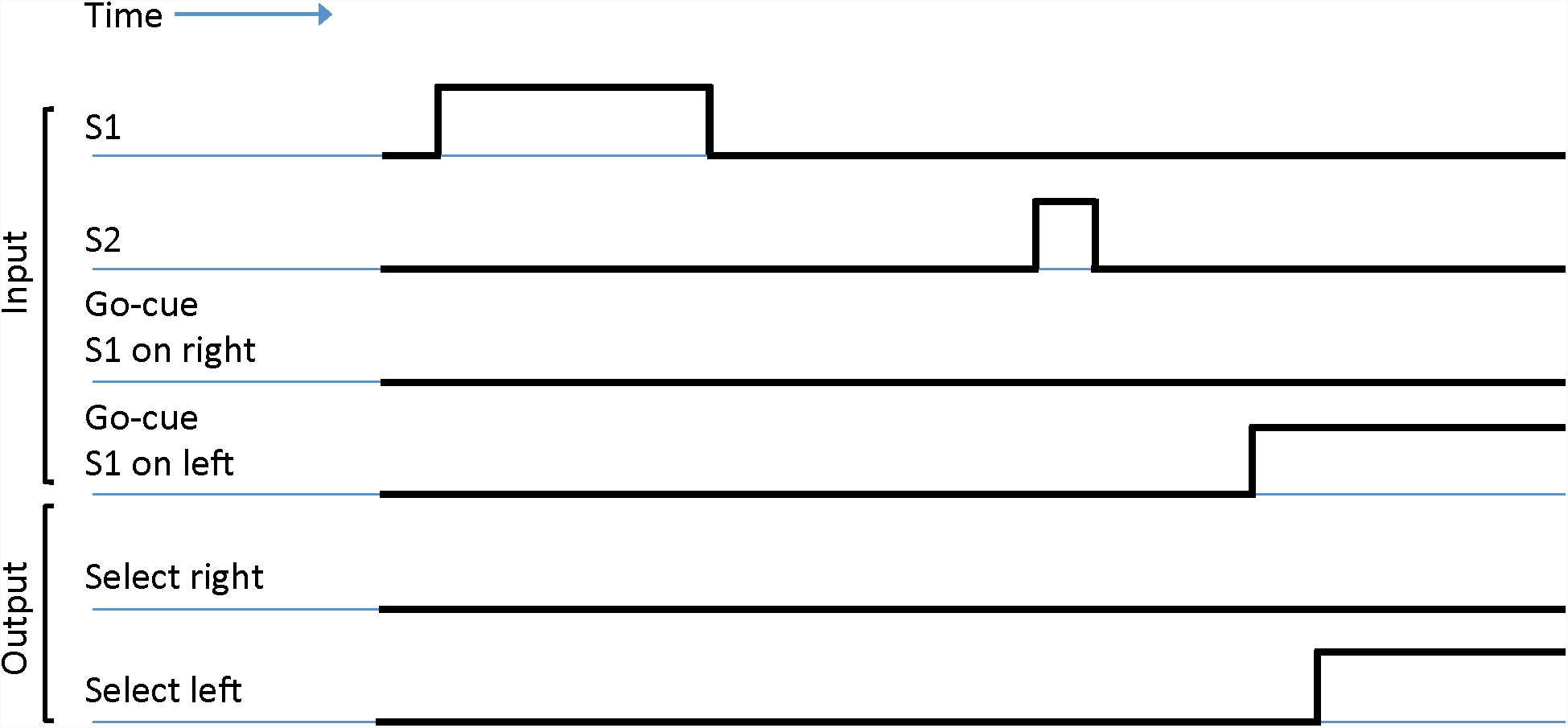
RNN inputs and outputs for the duration-discrimination task of Genovesio et al.^30^. The RNN compares the duration of two stimuli (S1 and S2) and then reports which stimulus was on longer.

### Stabilized random RNN

We analyzed the pretrained network accompanying the paper of Laje and Buonomano^10^. To ensure the number of units is similar across datasets, we randomly selected 100 out of the 800 units in the network for further analyses. We generated and analyzed 100 trials from the network by adding Gaussian random noise, with zero mean and standard deviation of 0.2, at each timestep of the simulation. The firing rate of each of the 100 units is stored every 100 ms and this activity is used in all subsequent analyses.

### Neural data

Electrode recordings were from nonhuman primates as previously described^15, 22, 29, 30^. For all datasets we only included correct trials in our analyses.

Context dependent working memory task of Saez et al.^15^: We analyzed units recorded for at least 50 trials in each of the four experimental conditions (context 1 or 2 and stimulus A or B) leading to 138 units in the Amygdala, 129 units in the OFC, and 102 units in the ACC.

Vibrotactile discrimination task of Romo et al.^22^: 160 PFC units were analyzed for the three second delay interval. Each unit was recorded for at least 10 trials for each value of f1 in the set [10 14 18 22 26 30 34] Hz. 139 PFC units were analyzed for the six second delay period. Each unit was recorded for at least 5 trials for each value of f1 in the set [10 14 18 22 26 30 34] Hz.

Ready-set-go interval reproduction task of Jazayeri and Shadlen^29^: 48 units were analyzed in LIP. Each unit was recorded for at least 20 trials with a minimum sample duration (interval between ready and set cues) of 1000 ms.

Duration-discrimination task of Genovesio et al.^30^: 148 units were analyzed in PFC. Each unit was recorded for at least 50 trials with a minimum S1 duration of 1000 ms.

## Supplementary Materials

**Figure S1:**
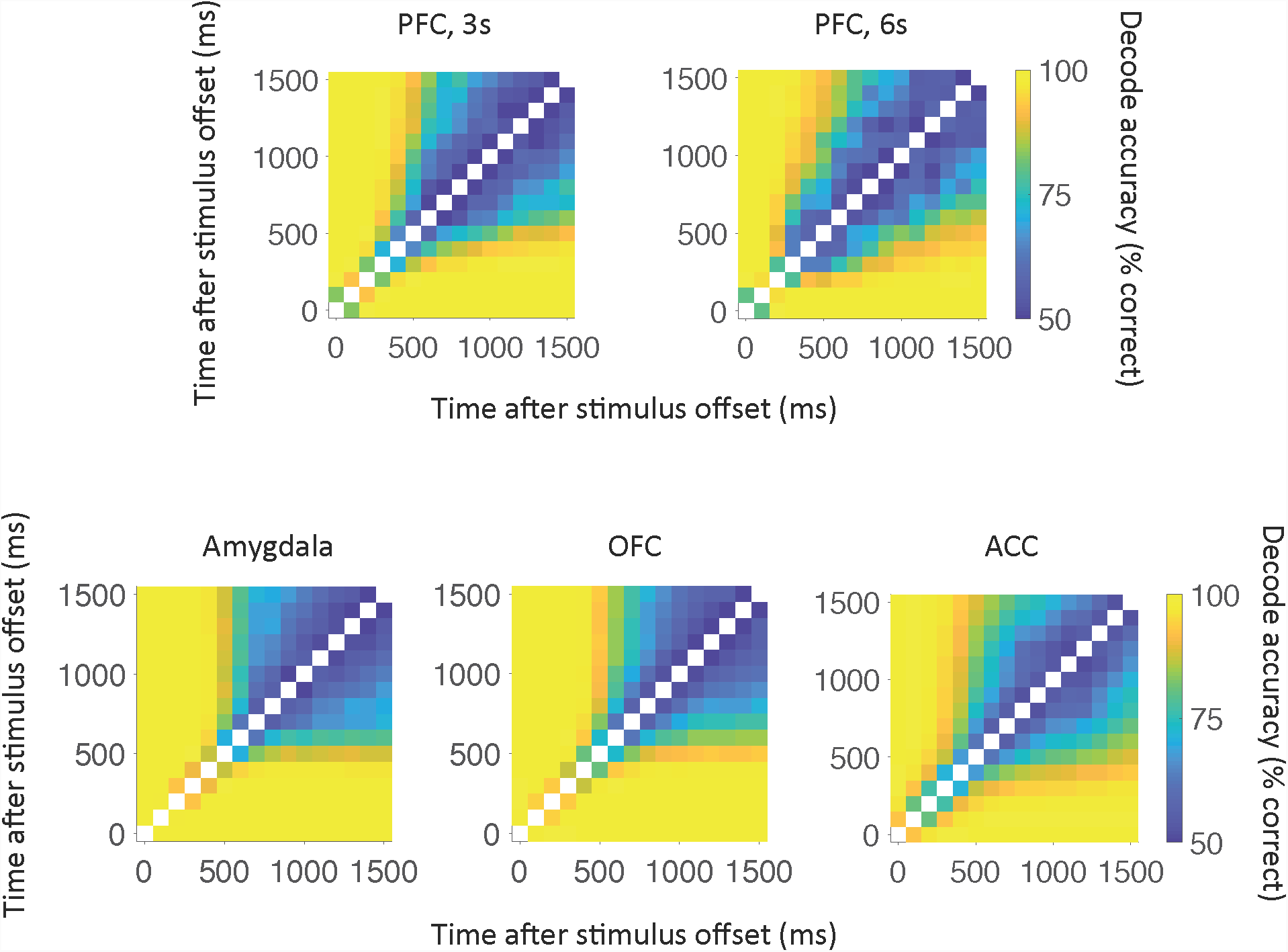
Two-interval time decode matrices for datasets of Romo et al.^22^ (top row) and Saez et al.^15^ (bottom row) during the 1500 ms interval after stimulus offset.

**Figure S2:**
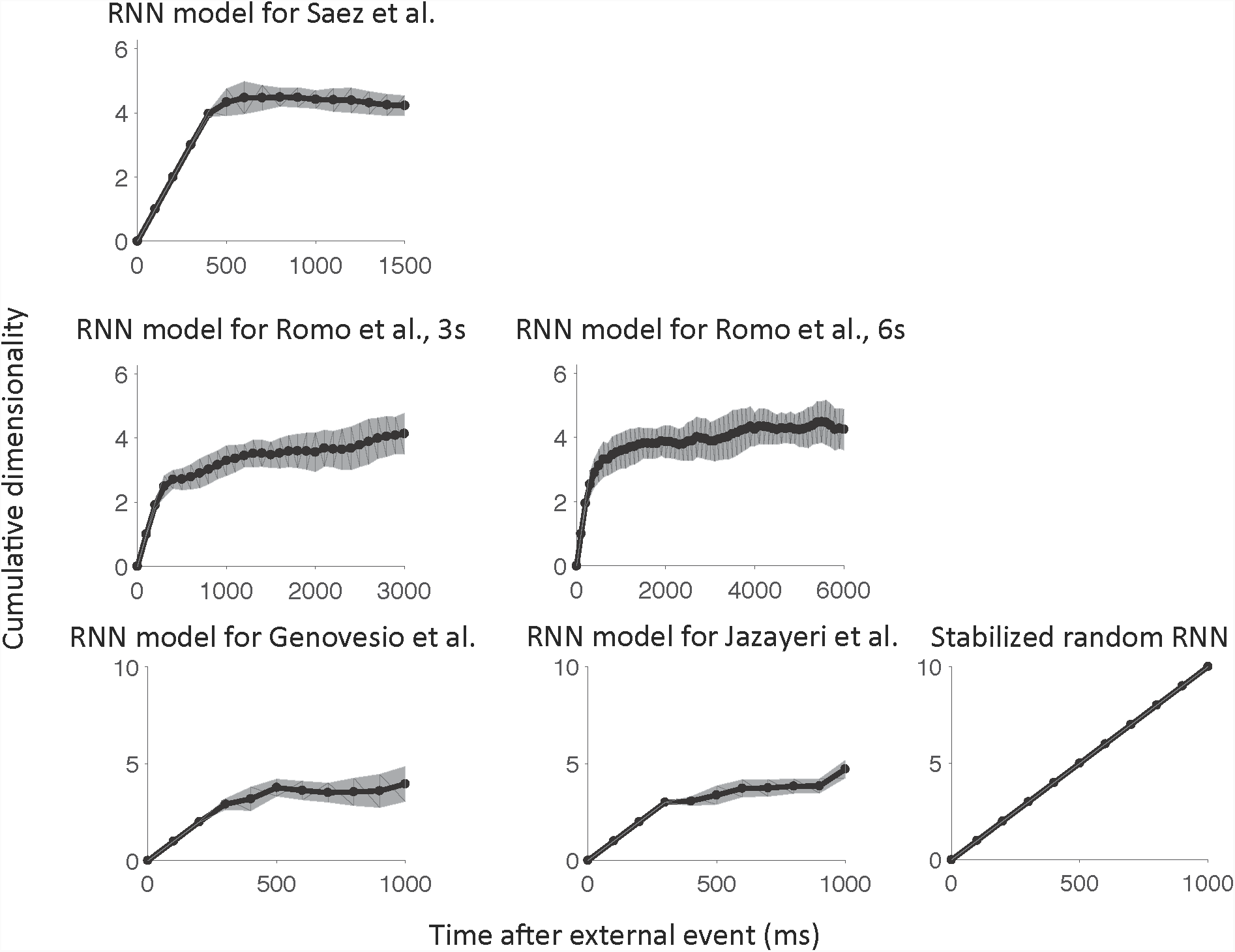
Cumulative dimensionality for RNN models. Error bars show one standard deviation.

**Figure S3:**
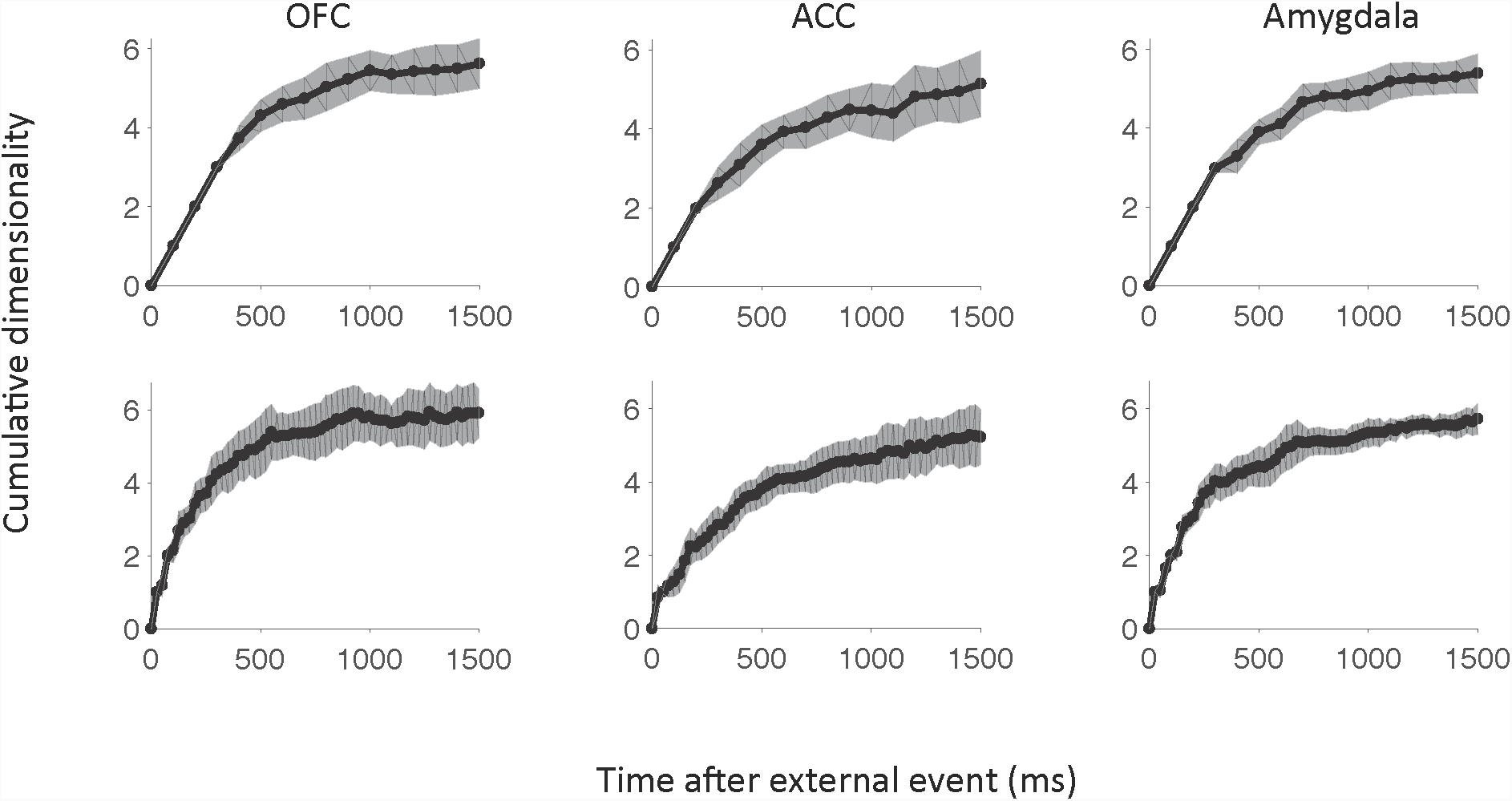
The cumulative dimensionality is stable as the number of timepoints along the neural trajectory are varied. In the top row, firing rates are from nonoverlapping 100 ms bins calculated every 100 ms. In the bottom row, firing rates are from overlapping 100 ms bins calculated every 25 ms. Data are from Saez et al.^15^ Error bars show one standard deviation.

**Figure S4:**
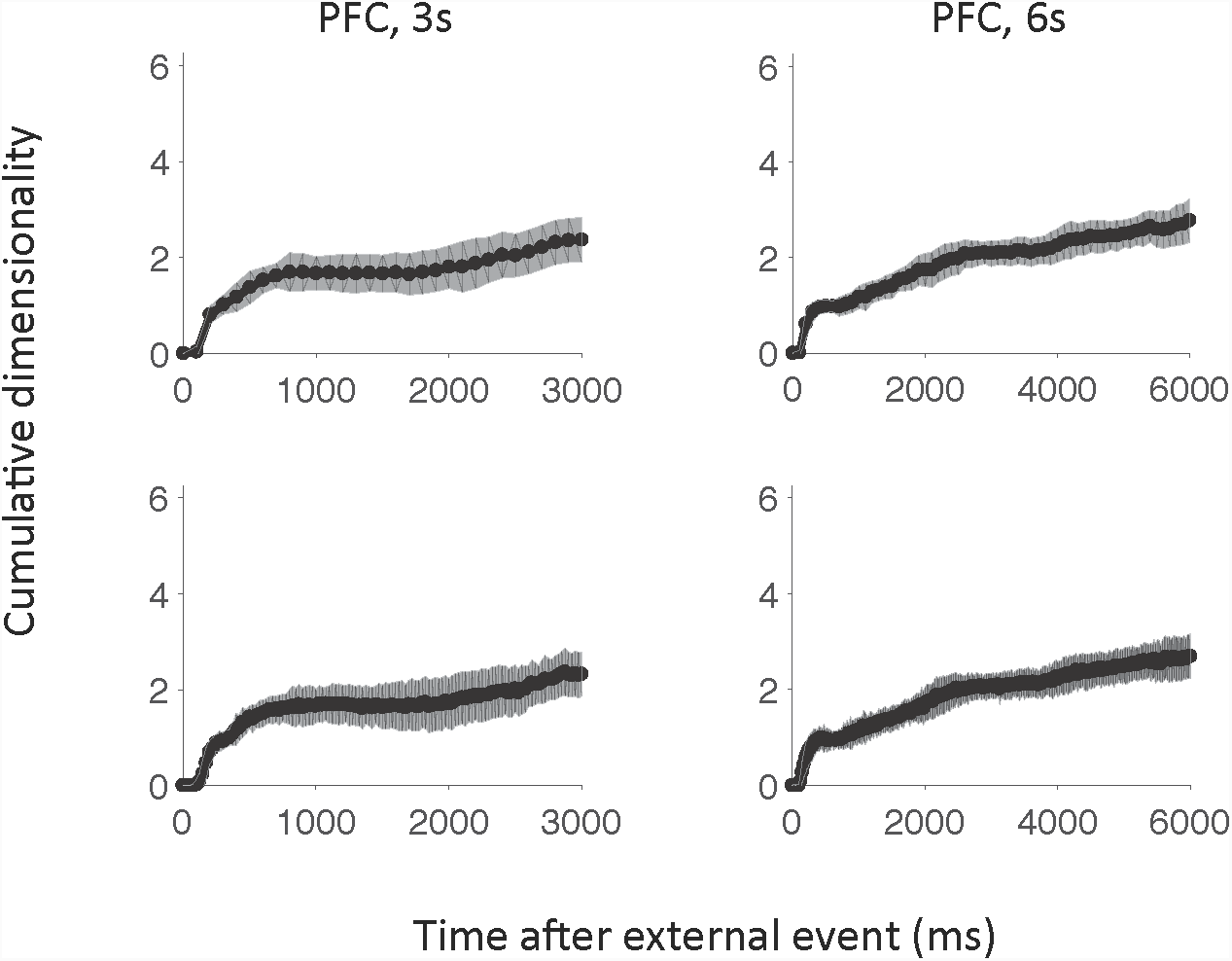
The cumulative dimensionality is stable as the number of timepoints along the neural trajectory are varied. In the top row, firing rates are from nonoverlapping 100 ms bins calculated every 100 ms. In the bottom row, firing rates are from overlapping 100 ms bins calculated every 25 ms. Data are from Romo et al.^22^ Error bars show one standard deviation.

**Figure S5:**
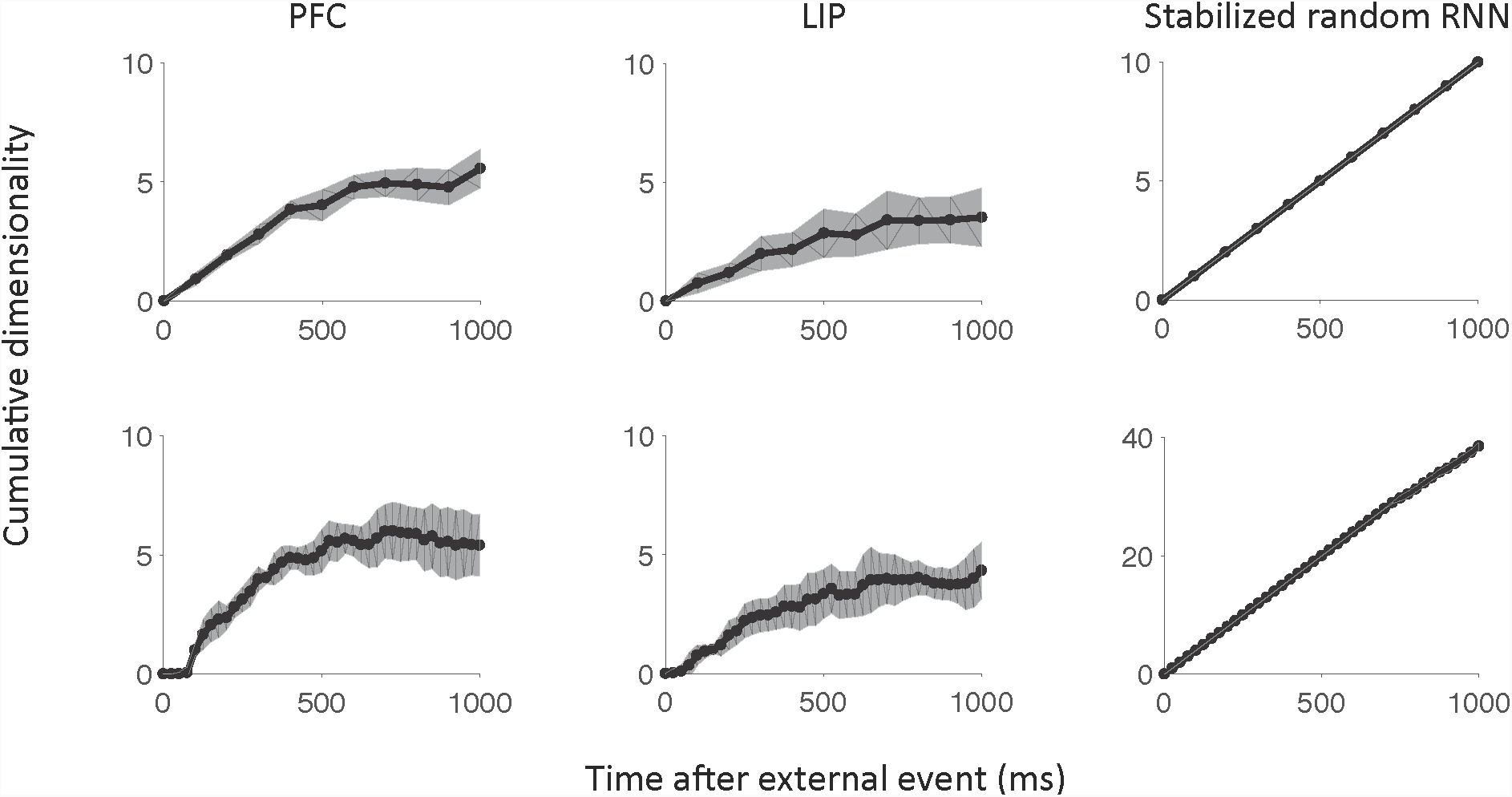
For the datasets of Genovesio et al.^30^ and Jazayeri et al.^29^, the cumulative dimensionality is stable as the number of timepoints along the neural trajectory are varied. In the top row, firing rates are from nonoverlapping 100 ms bins calculated every 100 ms. In the bottom row, firing rates are from overlapping 100 ms bins calculated every 25 ms. For the stabilized reservoir network of Laje and Buonomano^10^, the cumulative dimensionality grows as the number of timepoints are sampled more densely. Error bars show one standard deviation.

